# Pan-microbiome analysis along the human respiratory axis reveals an ecological continuum in health and collapse in disease

**DOI:** 10.1101/2025.09.04.674177

**Authors:** Tejus Shinde, Christina Kumpitsch, Yancong Zhang, Eric A. Franzosa, Rokhsareh Mohammadzadeh, Viktoria Weinberger, Tomas Mikal Eagan, Curtis Huttenhower, Vasile Foris, Christine Moissl-Eichinger

**Affiliations:** Diagnostic and Research Institute of Hygiene, Microbiology and Environmental Medicine, Medical University of Graz, 8010 Graz, Austria; Division of Respiratory Medicine, Department of Internal Medicine, Lung Research Cluster, Medical University of Graz, 8010 Graz, Austria; Department of Biostatistics, Harvard T.H. Chan School of Public Health, 02115 Boston, USA; Shenzhen Branch, Guangdong Laboratory of Lingnan Modern Agriculture, Genome Analysis Laboratory of the Ministry of Agriculture and Rural Affairs, Agricultural Genomics Institute at Shenzhen, Chinese Academy of Agricultural Sciences, Shenzhen, China; Department of Thoracic Medicine, Haukeland University Hospital, 5021 Bergen, Norway; Channing Division of Network Medicine, Department of Medicine, Brigham and Women’s Hospital, Harvard Medical School, 02115 Boston, USA; Broad Institute of Harvard and MIT, Cambridge, Massachusetts (MA); Department of Immunology and Infectious Diseases, Harvard T. H. Chan School of Public Health, Boston, MA; Harvard Chan Microbiome in Public Health Center, Harvard T. H. Chan School of Public Health, Boston, MA; BioTechMed-Graz, 8010 Graz, Austria

## Abstract

The human respiratory tract (RT) harbors complex microbial communities whose functions are critical to health and disease. Yet, current insights remain fragmented across anatomical sites, populations, and clinical states, limiting the field’s ability to define common patterns in health and disease.

Here, we present the first global respiratory pan-microbiome atlas, a resource integrating over 4,000 metagenomes across upper, intermediate, and lower RT from diverse cohorts encompassing health, pneumonia, COVID-19, and cystic fibrosis. Standardized taxonomic profiling reveals marked biogeographic structure: in health, lower RT communities largely represent filtered subsets of upper RT microbiota. Respiratory disease disrupts this organization, with reproducible depletion of core taxa at specific locations such as *Rothia mucilaginosa* and *Fusobacterium pseudoperiodonticum*, the latter being present in 88% of healthy sputum samples. Source-tracking analyses further support the collapse of inter-compartmental connectivity in disease and show the emergence of invasive taxa of unclear origin. Finally, prevalence-based models outperform abundance-based models in detecting disease-associated disruptions, providing greater sensitivity to shifts in community stability.

Altogether, this atlas defines the healthy RT microbiome as a spatially structured ecosystem and provides a foundational reference for advancing personalized care and systems-level models of respiratory disease.

## Introduction

The human respiratory tract (RT) represents one of the most extensive mucosal surfaces in the body, continuously exposed to airborne microbes and environmental particulates, with over 7000 litres of air and about 10^4^-10^6^ microbial cells inhaled every day^1^.

Unlike the unidirectional gastrointestinal tract, the RT has a branching architecture with bidirectional airflow, creating distinct ecological constraints on microbial persistence. Air moves bidirectionally through inhalation and exhalation, introducing microbes into the lower respiratory tract via inhalation and microaspiration^2^. Under physiologic conditions microorganisms reaching the distal lung cannot pass through; instead, their persistence is tightly constrained by elimination mechanisms such as mucociliary clearance, coughing, epithelial defenses, and immune surveillance- ecological pressures that contrast sharply with the unidirectional, continuous flow of the gut^1^. Unlike the GIT, where microbes encounter peristalsis and digestion, the RT presents a less permissive but highly dynamic environment, shaping microbial assembly and host interactions.

The RT is traditionally divided into the upper respiratory tract (URT) , including the anterior nares, nasal cavity, sinuses, nasopharynx, Eustachian tube, middle ear cavity, oral cavity, oropharynx, and larynx, and the lower respiratory tract (LRT) including the part of the larynx below the vocal cords, the trachea, bronchi and smaller airways of the lung^3–5^. In line with prior studies examining anatomical and microbial gradients across the respiratory tract, we refer to sampling sites such as the tongue dorsum, palatine tonsils, supraglottic area, throat, as intermediate respiratory tract (IRT) sites including sputum as a source of mixed origin. These regions are anatomically situated between the upper and lower respiratory compartments and are known to harbor microbial communities that reflect both oral and pulmonary origins ^6–8^.

Far from being sterile, the entire RT is now recognized as a complex microbial habitat, colonized by diverse communities of bacteria, fungi, archaea, and viruses that engage in dynamic interactions with each other and with the host^9–11^. These microbial populations are implicated in shaping immune responses, maintaining epithelial integrity, and modulating susceptibility to pathogens ^5,12,13^. Disruptions in RT microbiota - whether through host factors, antibiotic use, environmental exposures, or infectious agents - have been associated with a range of diseases, from acute lower respiratory infections to chronic airway disorders such as asthma, chronic obstructive pulmonary disease (COPD), cystic fibrosis (CF), and interstitial lung diseases ^7,9,14–20^, with the COVID-19 pandemic enhancing their clinical relevance^21^.

Despite the importance of RT microbiota, current knowledge remains fragmented. Most studies have focused on single diseases, specific regions or geographically restricted cohorts, limiting our ability to discern overarching principles of respiratory microbial ecology. Evidence suggests that microbial communities differ among sites such as the nasopharynx, oropharynx, and lower airways^4,6,7^, but whether these differences are consistent across individuals, populations, and disease states remains poorly defined. Moreover, the RT is a continuous, functionally integrated system, yet how communities are connected across compartments or how disruptions in one site influence others is underinvestigated. This piecemeal approach has limited our ability to discern overarching patterns of microbial biogeography and dynamics along the RT.

This lack of integrative analyses leaves critical questions unanswered: What is the baseline structure and variability of the respiratory microbiome along the RT in health? How do anatomical features, sampling sites, or host health states influence microbial assembly? How does the microbial flow across compartments, such as those from the upper or intermediate airways into the distal lung, change in health and disease?

To address these gaps, we conducted a pan-microbiome analysis of the human RT, integrating over 4,000 metagenomes from diverse cohorts across upper (anterior nares, nasal and oral sites, saliva), intermediate (tonsils, tongue dorsum, supraglottal, throat, sputum), and lower (BAL, lung tissue) compartments in health and disease. Using standardized taxonomic profiling (MetaPhlAn-4) and multivariate analyses, we delineated the spatial organization of respiratory communities and the relative influence of anatomy, sample type, and disease. We constructed a unified atlas of the respiratory microbiome in both health and disease, including pneumonia, COVID-19, and CF.

We show that respiratory microbial communities are strongly shaped by anatomical compartmentalization in health, but exhibit disrupted coherence in disease, with reproducible depletion of core taxa (e.g. *Fusobacterium pseudoperiodonticum*, *Prevotella melaninogenica*, and *Rothia mucilaginosa*) and altered microbial exchange across compartments. These findings redefine the RT as an interconnected microbial ecosystem whose continuity collapses in disease, and establish a pan-RT atlas as a foundational resource for diagnostics, therapeutics, and systems-level models of respiratory health advancing possibilities of personalized care.

## Results

### Broad respiratory microbiome dataset reveals variation in host–microbe ratios and major drivers of community structure

To capture a system-wide view of the microbial diversity across the human respiratory tract (RT), we assembled a large, cross-sectional collection of whole metagenome sequencing datasets from both public and author-contributed sources (N = 4,453 samples). These datasets covered samples from the upper, intermediate and lower RT (URT, IRT, and LRT), including nasal cavity (nasal swabs, anterior nares, nasopharyngeal aspirates), oral cavity (saliva, buccal mucosa, oral swabs, tongue dorsum), throat, sputum, and bronchoalveolar lavage (BAL), collectively spanning the full axis of the human RT. Anatomically, LRT metagenomes from sample types like BAL and lung tissue are moderate to highly invasive; therefore, we had fewer samples from the LRT compared to the URT sample types (URT: n=1,840 v/s LRT: n=872), where less invasive sampling strategies are more feasible (**Fig. 1-A**). The dataset spans healthy individuals and those with respiratory conditions, including pneumonia, cystic fibrosis, and COVID-19, capturing a representative spectrum of clinically relevant respiratory conditions associated with infections. Given the heterogeneity of data sources, we implemented a uniform preprocessing and analysis pipeline to enable consistent downstream comparisons (**Fig. 1-B**). This included automated download, metadata harmonization, quality control (QC), taxonomic profiling, and batch correction across studies (supplementary **Fig. S-1**).

**Figure 1:**
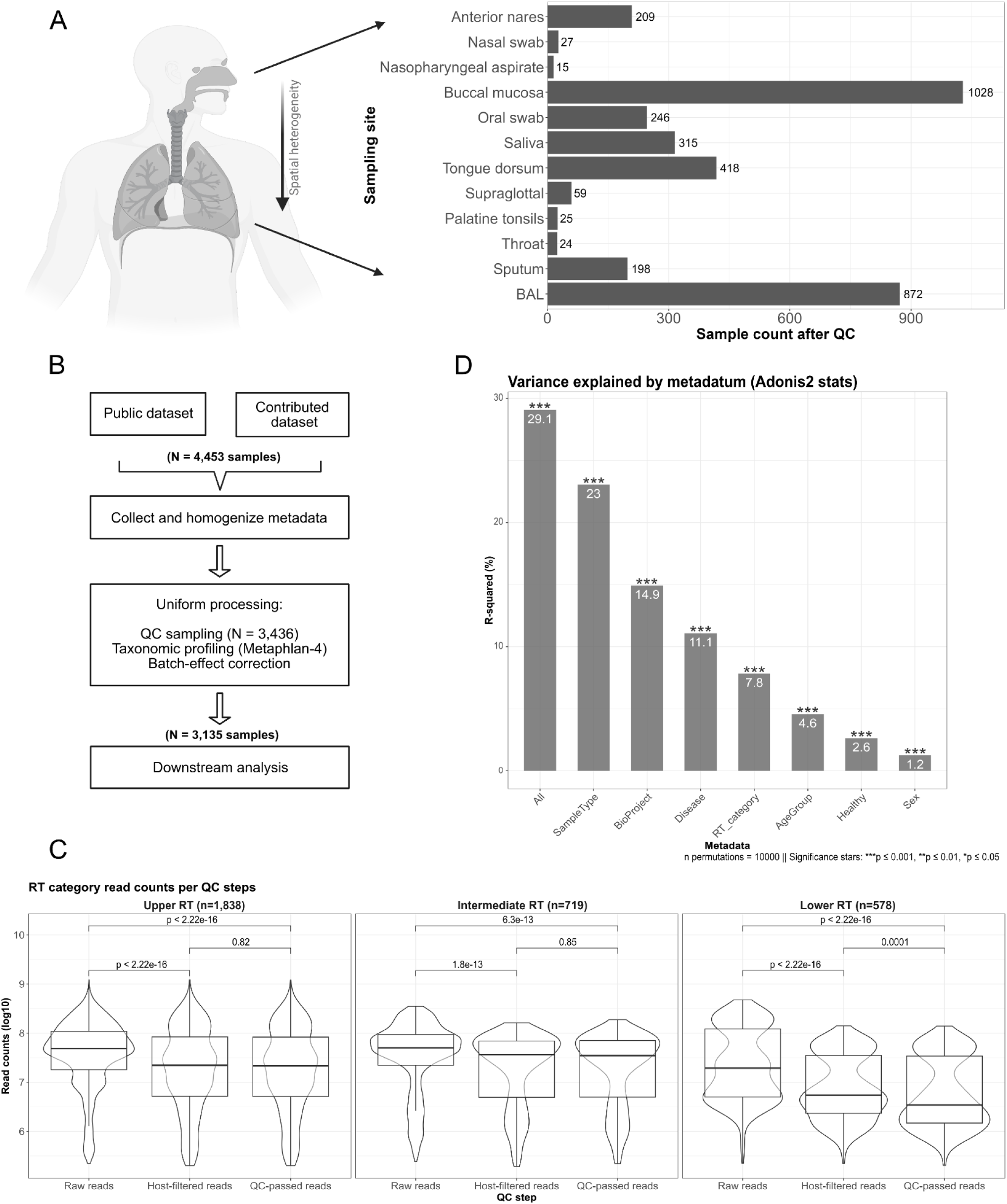
Respiratory microbiome dataset: anatomical distribution, processing pipeline, sequencing depth, and variance explained. (A) An anatomical diagram of the respiratory tract with sample distribution across body sites. (B) Schematic overview of the data processing pipeline for microbiome profiling and analysis used in this study. (C) Boxplot of sequencing read counts at different quality control stages (Raw, Decontaminated, Final QC). (D) Bar plot showing the univariate R-squared variance (post batch effect correction) explained by collected and homogenized metadata variables. All p-values in adonis2() were calculated using permutation tests (10,000 permutations).

Strict quality control (QC) of sequencing reads was critical, as respiratory microbiome samples are known to contain low microbial biomass, high host (human) DNA content, and exhibit lower diversity compared to gut microbiomes. To assess sequencing depth and sample retention across compartments, we tracked read counts across major QC stages. We analyzed more than 256 billion reads spanning the entire RT. After preprocessing, 3,135 high-quality metagenomic samples with a combined total of ∼165 billion reads were retained for further analysis. A detailed summary of read counts at each QC step, broken down by dataset, can be found in supplementary **Table S-1** & Supplementary **Fig. S-2**. The distribution of raw, host-filtered, and final QC-passed read counts for each RT category (URT, IRT, LRT) revealed substantial variability across sources. For instance, there was a significant difference in read counts before and after QC in all RT categories (Kruskal-Wallis group comparison between raw reads vs. QC-passed read counts; p < 0.005 for all RT categories) (**Fig. 1-C**). In particular, LRT samples lost a large proportion of reads following host decontamination, reflecting the high host-to-microbe ratio associated with invasive sampling, yet still retained sufficient microbial reads for analysis. Notably, our QC pipeline also filtered out low-biomass or degraded samples that could distort community structure or inflate the apparent abundance of rare taxa. In total, post-QC, 464 additional samples were excluded due to low-read counts, failed taxonomic profiling, or other filtering criteria, as detailed in the data processing flowchart (supplementary **Fig. S-1**).

Microbial communities can vary widely between individuals and sample types, but this variation is often shaped by underlying factors such as biogeography (e.g., nasal swab vs. lung fluid), disease status, age, and sex. Moreover, the study effect (reflecting differences in cohort composition, sampling protocols, and processing pipelines) can significantly influence observed microbial profiles.

To identify the major factors shaping respiratory microbiome variation in our data collection, we performed univariate PERMANOVA (adonis2 analysis on Bray–Curtis dissimilarity) for each metadata variable, both before and after batch (=study) effect correction. In the post-correction models, the *SampleType* and *BioProject* explained the largest proportions of variance (*R^2^* = 23. 0% & 14. 9% respectively), reflecting both anatomical sampling site and technical heterogeneity across studies (**Fig. 1-D**). Other factors, including specific disease labels (*Disease* variable; e.g., pneumonia, COVID-19), respiratory tract category (*RT_category*), age group (*AgeGroup* variable), health status (*Healthy* variable) and sex (*sex* variable), also showed statistically significant effects (permutation-based p ≤ 0.001; 10,000 permutations; **Fig. 1-D**), but with progressively smaller proportions of variation (R² =11.1 %; 7.8%; 4.6%; 2.6%; and 1.2%; respectively). In the multivariable model combining all variables, the cumulative explained variance reached 29.1%, indicating moderate but significant structuring of the respiratory microbiome by host health, sample type, and technical factors (**Fig. 1-D**). This suggests that while individual variables capture important variance components, much of their explanatory power overlaps, particularly between *SampleType*, *Disease*, and *BioProject*. Using batch effect correction the variance contributed by the *BioProject* variable dropped from 17.9% to 14.9%, suggesting a reduction in technical study-specific bias following correction (**Figs. 1-D & S-3**). The unexplained residual variance (∼71%) likely reflects unmeasured environmental, host genetic, or technical influences, as well as inherent inter-individual microbiome variability.

In this study, Disease and SampleType were designated as primary variables of interest, forming the basis of differential modeling and source tracking analyses. Additional variables, such as *RT_category* served as secondary stratifiers to support anatomical interpretation, while *AgeGroup* and *sex*, which explained relatively little variance (4.6% and 1.2%, respectively), were considered less informative in the global context of this dataset.

### Spatial gradients and ecological heterogeneity shape the respiratory microbiome

To investigate the major determinants of respiratory microbiome variation, we analyzed broad compositional patterns across sample types and respiratory tract regions. Principal coordinate analysis (PCoA) of Bray–Curtis dissimilarities revealed clear gradients in microbial composition by both *RT_category* and *SampleType* (**Fig. 2A–B**). *SampleType*, reflecting both anatomical origin and collection method (e.g., oral swabs or saliva to sample URT), emerged as a key driver of microbial structuring. A striking pattern was observed for anterior nares samples, which formed a tightly clustered group along the first principal coordinate (Supplementary **Fig. S-4**), indicating a highly constrained and compositionally distinct community. Given their dominant influence on β-diversity structure, anterior nares samples were excluded from downstream analyses to prevent confounding of broader ecological patterns. Re-ordination after removing anterior nares samples (**Fig. 2-B** & **Fig. S-5**) led to a marked shift in ordination geometry, enhancing the resolution of biologically coherent groupings. The remaining samples formed a broad continuum, with oral and other IRT sample types (e.g., saliva, tongue dorsum, and sputum) distributed between buccal mucosa and BAL, suggesting a smooth anatomical and ecological transition from upper to lower respiratory compartments. Notably, LRT (BAL) and IRT samples (sputum and tongue dorsum) partially overlapped in ordination space, reflecting proximity in β-diversity structure (**Fig. 2-B**).

**Figure 2:**
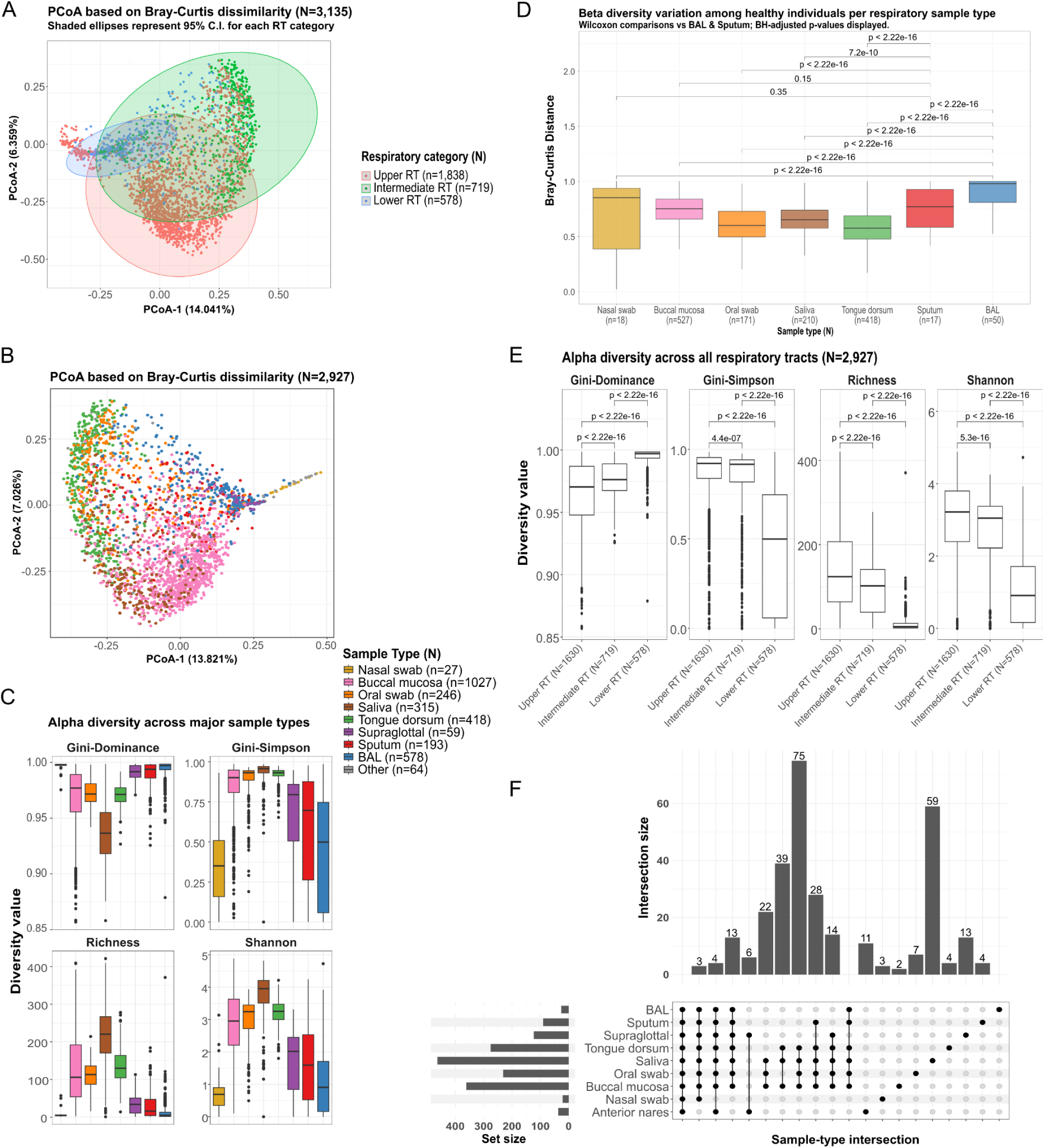
Microbial diversity, similarities and variance partitioning along the human RT. (A) Bray–Curtis β-diversity ordination (PCoA), colored by RT category with 95% confidence ellipses, reveals clustering and partial overlap among respiratory compartments. (B) Re-ordination after filtering out anterior nares and outlier samples shows that community structure clusters primarily by sample type. BAL and IRT samples (sputum and tongue dorsum) partially overlap in ordination space, reflecting proximity in β-diversity structure, whereas nasal swabs remained distinct with no overlap with other sample types. (C) Alpha diversity across respiratory tract sample types, measured using dominance (Gini), evenness (Gini–Simpson), richness, and Shannon index. Oral habitats (saliva, buccal mucosa) consistently exhibited higher richness and Shannon diversity compared to nasal and lower respiratory tract samples (BAL, sputum). (D) Bray–Curtis β-diversity among healthy individuals by sample type (colors as in C) shows that BAL communities are the most distinct, followed by nasal swabs and sputum, whereas oral habitats are more conserved across individuals. Although α- and β-diversity are not directly comparable, together these results highlight greater structural variability in nasal and BAL samples compared to oral sites. (E) Alpha diversity metrics aggregated by respiratory tract region reveal a clear gradient of decreasing richness and evenness from the URT to the LRT, reflecting a shift toward less diverse and more dominance-structured communities in the lower airways. (F) UpSet plot showing the number of microbial species shared across major respiratory tract sample types (minimum 10-sample prevalence threshold). Most species were shared across multiple compartments, supporting ecological connectivity along the respiratory tract. By contrast, site-specific taxa were rare (e.g., anterior nares), consistent with their distinct and compositionally constrained profiles.

Species accumulation curves across respiratory tract compartments (Supplementary **Fig. S-6A**) highlight both the scale and the novel contribution of our multi-cohort dataset. LRT and IRT categories showed early plateauing trends, suggesting that species richness in these compartments is closer to saturation. By contrast, the URT curve continued to rise, largely reflecting its larger sample size and greater microbial diversity. Similar patterns were observed at finer resolution (Supplementary **Fig. S-6B**), where buccal mucosa and oral swabs displayed especially high species accrual, underscoring the central role of oral niches in driving overall respiratory tract diversity.

To quantify microbial diversity across the respiratory tract, we examined within-sample (alpha) and between-sample (beta) diversity in healthy individuals (**Fig. 2-C & 2-D**). Alpha diversity analysis revealed strong, compartment-specific ecological patterns (**Fig. 2-C**). Nasal swabs and BAL samples (representing anatomical extremes of the URT and LRT) consistently showed the lowest species richness and Shannon diversity, alongside elevated Gini-dominance values, indicating that these low-diversity locations are dominated by a few taxa. In contrast, oral cavity-associated sites (e.g., saliva, buccal mucosa) and IRT samples (e.g., sputum) displayed higher richness and evenness, suggesting more complex and balanced microbial communities. Collectively, these trends revealed an inverted U-shaped gradient in diversity across the respiratory axis, peaking in oral and IRT sites and tapering off toward the URT and LRT ends.

While alpha diversity captures the internal complexity of each sample, beta diversity sheds light on how microbial communities vary across individuals within each site. Beta diversity analysis highlighted this inter-individual variation (**Fig. 2-D**), with nasal and BAL samples exhibiting the highest beta diversity, indicating considerable heterogeneity between individuals. In comparison, saliva, tongue dorsum, and sputum samples showed significantly lower beta diversity, suggesting a more conserved microbial community structure across individuals. These opposing diversity patterns (high alpha and low beta diversity in oral sites versus low alpha and high beta diversity in nasal and BAL samples) suggest that different regions of the respiratory tract are shaped by distinct ecological forces: oral sites may host stable, diverse communities common across individuals, while nasal and lower airway niches support more individualized, lower-diversity communities likely influenced by stronger environmental filtering, spatial isolation, or host immune pressure. Our finding that saliva harbors a rich, even, yet relatively conserved microbiota is consistent with observations from the Human Microbiome Project (HMP), which characterized oral microbial communities in healthy individuals^22^. In contrast, the compositional constraint and high inter-individual variability of BAL and nasal samples suggest that distal respiratory sites may be more influenced by stochastic colonization or localized and specific host factors.

Stratification of samples by RT category (URT, IRT, and LRT) reinforced the picture of an anatomical gradient (**Fig. 2-E**). LRT samples (BAL) exhibited the lowest richness and evenness (reflected in significantly reduced Shannon diversity and elevated dominance indices), known to be the hallmarks of compositionally constrained niches (BH*-*adjusted p< 2.22 × 10⁻¹⁶ for LRT v/s URT and LRT v/s IRT). URT samples showed the highest alpha diversity and evenness, while IRT samples (saliva, sputum, tongue dorsum) consistently fell in between, reflecting their transitional anatomical location. This diversity gradient supports the concept of the respiratory tract as a continuous ecological landscape rather than a series of discrete compartments.

#### Species are shared from the upper to the lower respiratory tract

To explore species overlap across respiratory sites, we constructed an UpSet plot of shared taxa across sample types (**Fig. 2-F**). The largest intersection comprised 75 species present across multiple URT sites (saliva, tongue dorsum, buccal mucosa, oral swab). Additional intersections of intermediate size (13–39 species) linked URT, IRT (sputum, supraglottal, tongue dorsum), and LRT (BAL) combinations, further highlighting a continuum of microbial connectivity along the respiratory axis. By contrast, site-specific taxa were comparatively rare: while no species were unique to BAL, 13 were restricted to anterior nares, consistent with the distinct and compositionally constrained nature of this niche. Nasal and anterior nares samples contributed relatively few shared taxa with other regions, consistent with their constrained and distinct phylum-level profiles. Together, these patterns suggest that while the respiratory tract harbors localized microbial subsets, the majority of taxa are shared across multiple compartments, supporting a model of downward microbial flow from oral reservoirs into the distal lung.

### The healthy respiratory microbiome exhibits spatial continuity and compositional filtering across compartments

Building on these diversity patterns, we next examined the taxonomic composition and distribution of microbial lineages across healthy respiratory compartments to assess community continuity and ecological filtering. To broadly characterize the taxonomic composition of healthy respiratory tract samples, we initially quantified the relative abundance and prevalence of microbial taxa at the domain level across sample types (**Fig. 3-A & Fig. S-7**). As expected, bacteria dominated the microbial landscape across all respiratory sites, with consistently high abundances in oral cavity-associated sites such as buccal mucosa, saliva, and tongue dorsum. In contrast, eukaryotic reads were markedly lower overall, though detectable in some sample types (particularly buccal mucosa and oral swabs), likely reflecting fungal signatures. Archaeal sequences were detected only at trace levels in the URT samples (particularly in buccal mucosa), and were nearly absent in intermediate and lower tract sites. While methanogenic archaea such as *Methanobrevibacter oralis* are well-documented in subgingival niches and have been associated with human periodontitis^23^, their restricted detection in our dataset suggests limited dispersal or colonization beyond the oral cavity into the lower respiratory tract. The trends confirm bacteria as the predominant microbial domain in the healthy respiratory ecosystem and reveal subtle site-specific variation on the domain level. (**Fig. S-7**).

**Figure 3:**
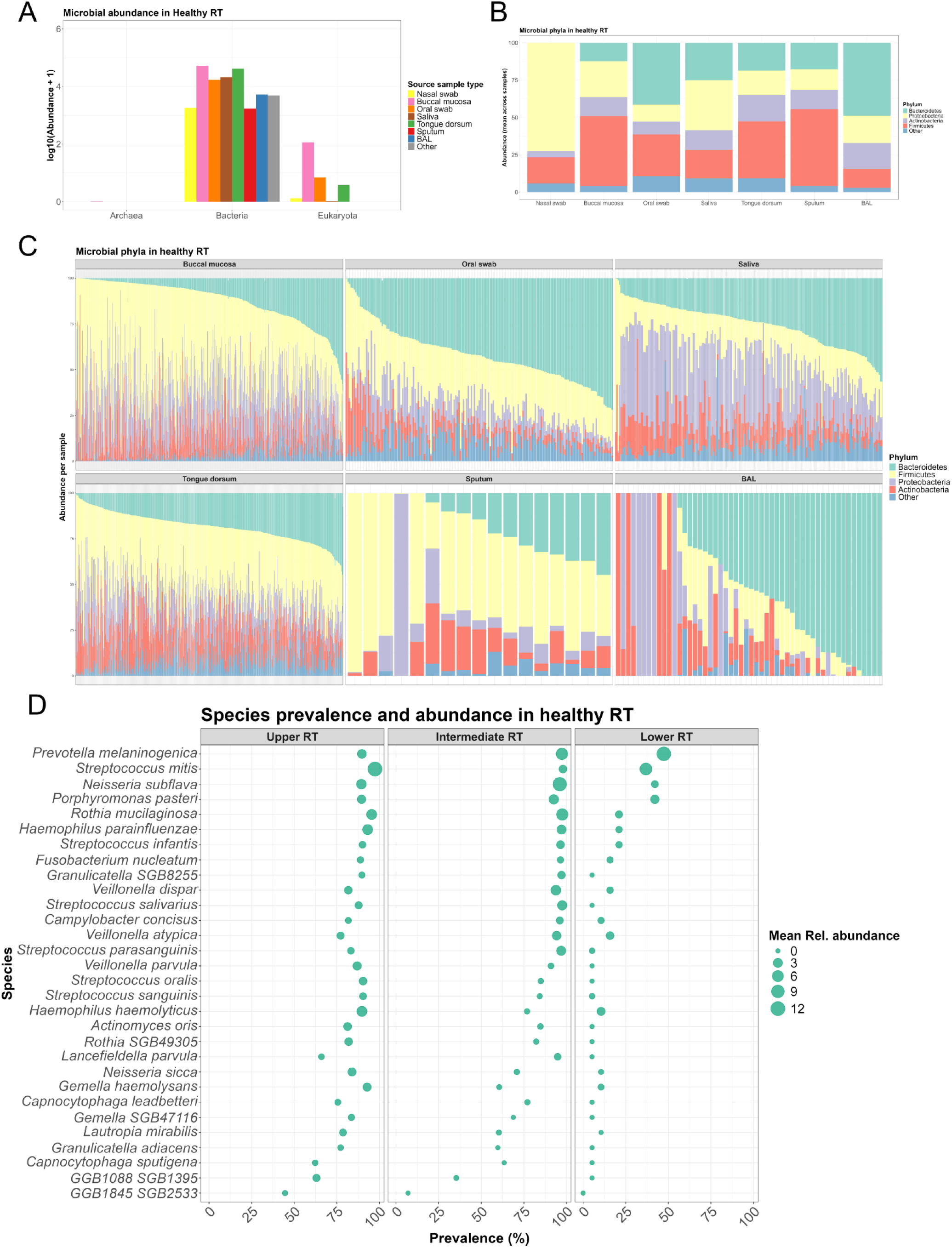
Microbial taxonomic composition of the ‘Healthy’ human RT across anatomical compartments. (A) Domain-level microbial abundance across respiratory sample types in healthy individuals shows bacterial dominance across all sites, with minimal archaeal and variable eukaryotic signals. (B) Mean phylum-level composition across sample types reveals a shift in dominant lineages from Bacteroidetes, Firmicutes, Proteobacteria, and Actinobacteria in the upper tract to increased Proteobacteria and Bacteroidetes in lower sites such as sputum and BAL, respectively. (C) Barplot for taxonomic composition from major sample types sorted by Bacteroidetes abundance. (D) Prevalence and mean relative abundance of the top 25 most prevalent species across the upper, intermediate, and lower respiratory tract. While several core species (e.g., *Prevotella melaninogenica*, *Rothia mucilaginosa*, *Streptococcus mitis*) are consistently present across compartments, others show sharp gradients in prevalence, suggesting spatial filtering or niche specificity.

Phylum-level profiles revealed pronounced compositional shifts across respiratory sites (**Fig. 3-B**). Nasal swabs were overwhelmingly dominated by Proteobacteria, in stark contrast to oral and BAL samples, where Bacteroidetes and Firmicutes predominated. Oral-associated URT samples (buccal mucosa, oral swabs, saliva) were characterized by a mixed dominance of Bacteroidetes and Firmicutes, with Actinobacteria contributing variably across individuals. In contrast, IRT-associated, tongue dorsum & sputum communities showed a strong predominance of Firmicutes, while BAL samples were consistently enriched for Bacteroidetes, accompanied by lower yet notable contributions from Firmicutes, Proteobacteria, and Actinobacteria. To capture inter-individual variation in phylum-level composition, we examined stacked barplots of relative abundance across healthy individuals (**Fig. 3-C**). While overarching dominance patterns were clear, Bacteroidetes were enriched in oral and BAL samples, and Firmicutes in tongue dorsum & sputum. However, this visualization revealed substantial within-site variability, particularly in URT and IRT sites, where no single phylum consistently prevailed across individuals. In contrast, BAL samples displayed a more uniform enrichment of Bacteroidetes, alongside notable but lower abundances of Proteobacteria, Actinobacteria, and Firmicutes, suggesting tighter ecological filtering in the lower airways.

To explore species-level variation across the respiratory tract, we examined the prevalence and relative abundance of the top 25 most prevalent species in healthy samples (**Fig. 3-D**). The URT (represented by nasal swabs, oral swabs, buccal mucosa, and saliva) was consistently dominated by commensals such as *Streptococcus mitis*, *Neisseria subflava*, *Haemophilus parainfluenzae*, *Prevotella melaninogenica*, and *Rothia mucilaginosa*, which were both highly prevalent and abundant (relative abundance). IRT samples, including sputum and supraglottic swabs, showed a broader taxonomic profile, with increased prevalence of *Veillonella*, *Actinomyces*, *Fusobacterium*, and *Prevotella* species. In the LRT, fewer species were detected overall, and these occurred at lower prevalence and abundance, with a subset of URT-and IRT-associated taxa (e.g., *Prevotella melaninogenica*, *Veillonella atypica*, *Streptococcus mitis*) still present. Additionally, a small number of low-prevalence but sporadically high-abundance species, such as Moraxella nonliquefaciens and Ralstonia pickettii, were observed in BAL samples (**Fig. S-8**).

### Pneumonia and COVID-19 are characterized by loss of core microbial species prevalence rather than abundance

To move beyond community-level differences in diversity and composition, we next pinpointed specific microbial taxa reproducibly associated with respiratory diseases, including pneumonia, COVID-19, and CF, across sample types and datasets.

To understand how respiratory diseases alter the lung microbiome, we focused on BAL samples for pneumonia and COVID-19, as these cohorts provided sufficient sample sizes in the lower respiratory tract, enabling robust statistical testing. Using MaAsLin3 modeling, we detected robust disease-specific associations at both the species prevalence and abundance levels (**Fig. 4**). The majority of statistically significant disease-linked shifts were driven by differences in species prevalence, rather than within-sample abundance, highlighting a loss of taxa across individuals, rather than changes in relative abundance (12 prevalence- v/s one abundance-based association for COVID-19, and 102 prevalence v/s one abundance based association, before multiple testing correction) (**Supplementary Table-4**).

**Figure 4:**
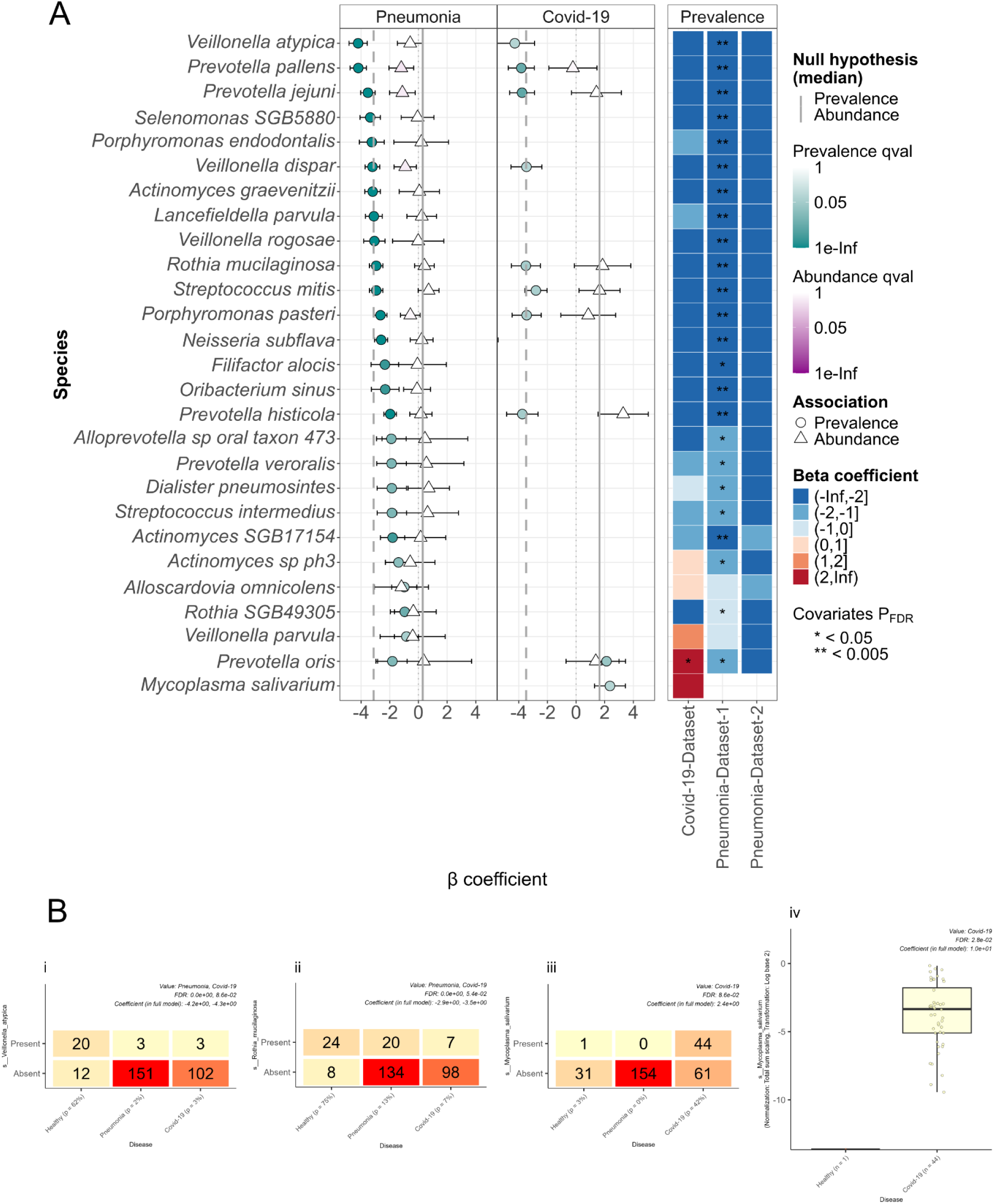
Disease-associated shifts in microbial prevalence and abundance in BAL (LRT) samples. (A) Differential species prevalence (circles) and abundance (triangles) in BAL samples were modeled using MaAsLin 3 on batch-corrected MetaPhlAn 4 profiles. Effect sizes (β-coefficients) with standard errors are shown on the left; negative values indicate depletion relative to healthy controls. Pneumonia was marked by widespread core taxa loss, including *Veillonella atypica*, *Prevotella pallens*, and *Rothia mucilaginosa*. The right panel shows β coefficients from dataset-specific models, with significance denoted by FDR-adjusted q < 0.05 (*) and q < 0.005 (**). Results highlight reproducible, prevalence-driven microbial disruptions in disease. Top 10 significant species per disease group and model type are shown. (B) Panels (i–iii) display contingency-style plots of species presence/absence across disease groups, based on MaAsLin 3 prevalence models. *V. atypica* (i) and *R. mucilaginosa* (ii) were significantly depleted in both pneumonia and COVID-19, while *Mycoplasma salivarium* (iii) was enriched in COVID-19. Panel (iv) shows the log-transformed, TSS-normalized relative abundance of *M. salivarium*, revealing a marked increase in COVID-19 samples (n = 44) compared to healthy controls (n = 1). Only one out of 32 healthy BAL samples harbored *M. salivarium*, versus 44 out of 105 in COVID-19, reinforcing the disease association and aligning with the positive β coefficients observed in both prevalence and abundance models. All panels report β coefficients and FDR q-values from the full MaAsLin 3 model. Together, these results highlight reproducible, disease-specific compositional shifts in the lower respiratory tract microbiome.

Across BAL samples, pneumonia was associated with 103 differentially prevalent or differentially abundant species (p-value < 0.05), including 102 prevalence-based and only 1 abundance-based association. Only the 100 prevalence-based species associations remained significant after multiple testing corrections (FDR q-value < 0.05). COVID-19 samples revealed 13 such associations (12 prevalence-based and one abundance-based). Out of these 13 species, only one uncultivated species (GGB1188_SGB1538*; β = 6.98,* q = 0.0277) was significantly associated with sequencing depth, suggesting limited technical confounding in the final models.

All 102 prevalence-based associations for pneumonia were negative, indicating widespread loss of taxa in that disease. In contrast, a few species, viz., *Prevotella oris* (prevalence association)*, Mycoplasma salivarium* (both prevalence and abundance model), exhibited a positive β-coefficient, i.e, showing increased abundance or prevalence in COVID-19. Among the most consistently depleted taxa were oral core species such as *Veillonella atypica*, *Prevotella pallens*, *Streptococcus infantis*, and *Actinomyces graevenitzii*, which showed significant loss in prevalence under disease conditions (q < 0.05 for all associations; **Fig. 4-A**). Core taxa such as *Rothia mucilaginosa*, *Streptococcus mitis*, and *Prevotella histicola* also declined across both diseases, indicating a cross-disease erosion of the health-associated lung microbiota. These prevalence-driven shifts are further illustrated in contingency-style plots of representative taxa (**Fig. 4-B**), highlighting the consistent depletion of core commensals and the enrichment of M. salivarium in COVID-19. A comprehensive list of the top 50 disease-associated taxa identified by MaAsLin3, including effect sizes and significance values, is provided in Supplementary **Fig. S-9**.

Only a handful of species exhibited strong abundance increases in disease. For example, *Haemophilus influenzae* (*β* = 5.50, q = 0.22) and *M. salivarium* (*β* = 9.97, q = 0.028) were more abundant in both pneumonia and COVID-19 cases, potentially reflecting opportunistic outgrowth in immunocompromised airways (Supplementary **Table-4**). However, such patterns were the exception rather than the rule.

To assess the consistency of species–disease associations across datasets, we complemented the cross-cohort model with separate prevalence and abundance modeling within each disease cohort (Model: ∼ *Disease* + *Total_reads*). This cohort-stratified approach enabled the detection of both study-specific effects and taxa that were reproducibly associated with disease. Several species, including *Rothia mucilaginosa* and *Prevotella histicola*, displayed consistently negative β-coefficients across all pneumonia and COVID-19 cohorts, suggesting robust and cohort-independent depletion of these commensals in disease (heatmap **Fig. 4-A)**. *Veillonella atypica* emerged as a particularly strong candidate for a generic loss-of-health biomarker, being significantly less prevalent in both pneumonia and COVID-19 groups in the primary model and consistently depleted across all disease datasets in the cohort-aware analysis.

Together, these findings reveal that respiratory diseases are marked by reproducible loss of commensal taxa, with prevalence-based models offering greater sensitivity to microbial disruption than abundance-based approaches. The depletion of core species across pneumonia and COVID-19 cohorts underscores a shared ecological signature of disease progression in the lower respiratory tract.

### Consistent depletion of core taxa in the sputum microbiota of cystic fibrosis patients

Following the analysis of pneumonia and COVID-19 in the lower respiratory tract, we next examined species-level alterations in the intermediate respiratory tract. For cystic fibrosis, sputum was selected as the primary comparator, given its status as the most abundantly represented sample type in this disease group and its clinical relevance as a proxy for lower airway microbial communities. Similar to LRT-associated diseases, prevalence shifts emerged as the dominant signal, with most species showing significant associations with CF based on presence/absence rather than relative abundance (69 prevalence-based vs. 3 abundance-based associations; **Supplementary Table-5**). The CF disease model identified 39 species with significant prevalence-based associations (p < 0.05), of which only two - *Leptotrichia wadei* (*β* = -3.51, q = 0.046) and *Herbaspirillum huttiense* (*β* = -5.95, q = 0.012) *-* remained significant after false discovery rate correction (q < 0.05) (**Fig. 5-A**). We highlight some of these prevalence-driven shifts with contingency-style plots of representative taxa in **Fig. 5-B**.

**Figure 5:**
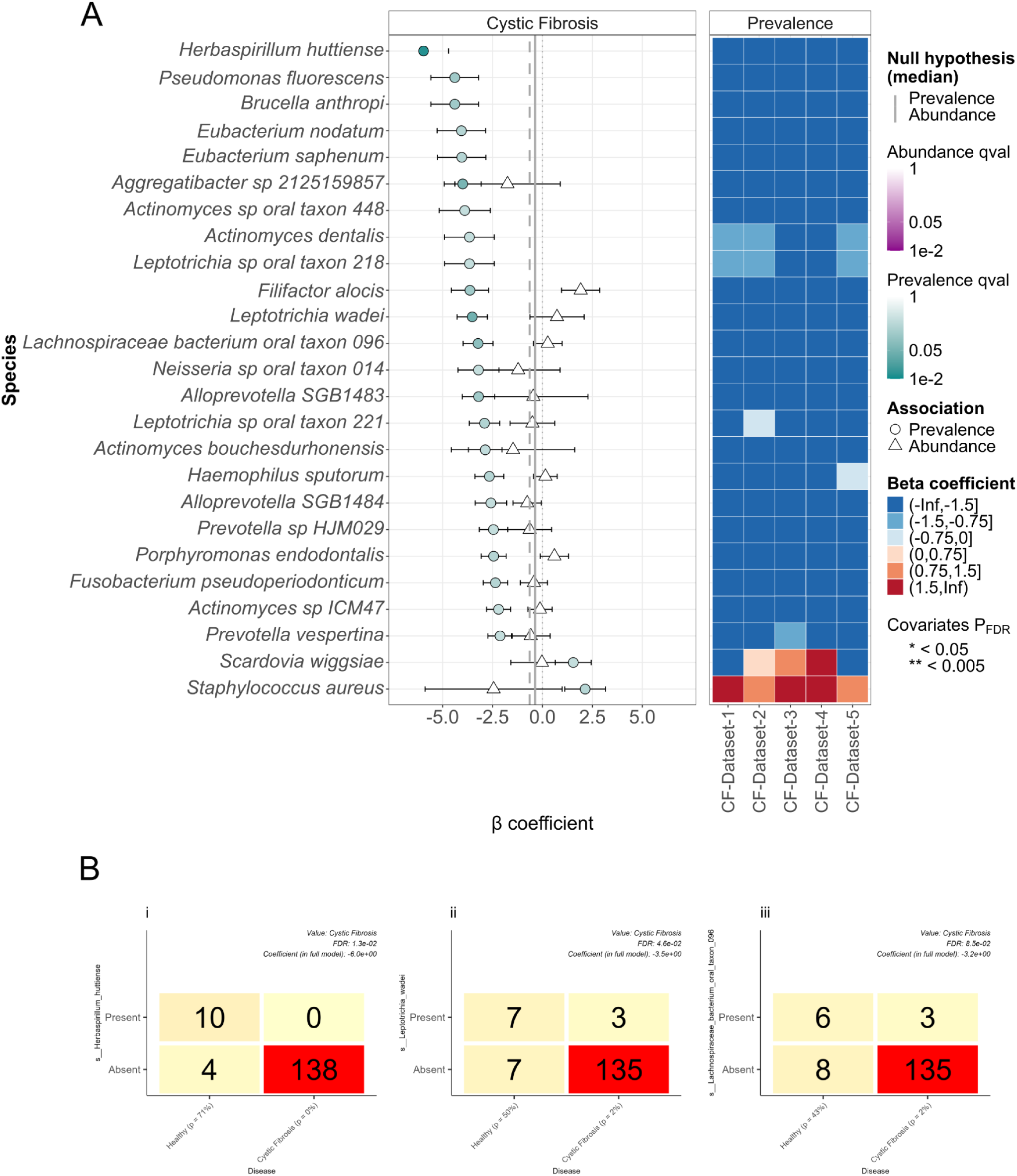
Disease-associated shifts in microbial prevalence and abundance in Sputum (IRT) samples. (A) Differential species prevalence (circles) and abundance (triangles) were modeled using MaAsLin 3 on batch-corrected MetaPhlAn 4 profiles. The left panel shows effect sizes (β coefficients) with standard errors; negative values indicate depletion relative to healthy controls. The right panel displays β coefficients from dataset-specific models, with statistical significance denoted as q < 0.05 (*) and q < 0.005 (**). Consistent prevalence loss patterns for species such as *Herbaspirillum huttiense*, *Leptotrichia wadei*, and *Lachnospiraceae bacterium* oral taxon-096 across all five CF datasets underscore their potential as robust, non–cohort-specific microbial indicators of cystic fibrosis. Only the top 10 species associations per disease group and model type are shown. (B) Panels (i–iii) show contingency-style plots of species presence/absence across disease groups, based on MaAsLin 3 prevalence models. *H. huttiense* (i), *L. wadei* (ii), and *Lachnospiraceae bacterium* oral taxon-096 (iii) were significantly depleted in CF samples. All panels report β coefficients and FDR-adjusted q-values from the full model. These findings highlight reproducible, disease-specific compositional disruptions in the respiratory tract microbiome.

A range of oral and anaerobic core species were depleted in CF, including *Peptostreptococcus stomatis, Filifactor alocis, Alloprevotella tannerae, Porphyromonas endodontalis*, and *Lachnospiraceae* oral-taxon-096. These organisms are commonly associated with healthy oral or upper airway niches^24^ and with lower airways^25^; their reduced prevalence in CF sputum suggests a broad loss of health-associated community members even in the IRT.

None of the 24 species associated with the sequencing depth variable (*Total_reads*) - 15 in the prevalence model and 9 in the abundance model (Supplementary Table-4) - were statistically significant after false discovery rate (FDR) correction at q < 0.05. We included sequencing depth as a covariate to identify and control for potential technical confounding; the lack of significant associations after correction suggests that disease-related signals were not driven by variation in sequencing depth.

Like LRT modeling done above, we also complemented the sputum cross-cohort model with separate prevalence and abundance modeling within each of the five disease cohorts (Model: ∼ 𝐷𝑖𝑠𝑒𝑎𝑠𝑒 + *Total_reads*). Consistent prevalence loss patterns (negative beta coefficients; for species like *Herbaspirillum huttiense, Pseudomonas fluorescens, Brucella anthropi*, *Eubacterium nodatum*, and *Eubacterium saphenum*) across all five CF datasets confirm that these prevalence associations are not cohort-specific. Although many of these species were not statistically significant in the cross-cohort model or individual cohort models, it can be argued that such consistent prevalence loss patterns across datasets strengthen their candidacy as robust CF-associated microbial markers.

Taken together, these findings suggest that CF-related disruption of the IRT microbiome is primarily characterized by changes in species presence or absence, rather than shifts in their relative abundance.

### Microbial source tracking reveals disrupted dispersal and untraceable microbial inputs in diseased respiratory niches

To complement our compositional modeling of the respiratory microbiome, we applied microbial source tracking to infer the origins of microbial communities in the lower and intermediate respiratory tract. Using BAL and sputum samples as proxies for the LRT and IRT, respectively, we assessed how disease influences microbial exchange along the respiratory tract.

In healthy individuals, BAL samples received a diverse set of microbial contributions from canonical upper and intermediate airway sources - including buccal mucosa, oral swabs, tongue dorsum, and saliva - reflecting intact dispersal and selection mechanisms along the respiratory axis (**Fig. 6-A**). By contrast, BAL samples from pneumonia and COVID-19 patients showed a striking reduction in these known source contributions. Instead, microbial communities were increasingly dominated by taxa of ’*Unknown*’ origin, suggesting colonization from uncharacterized reservoirs or environmental seeding pathways. Statistical comparisons confirmed that proportional contributions from buccal mucosa, oral swabs, tongue dorsum, and sputum were significantly reduced in both disease groups (Wilcoxon rank-sum test comparing healthy vs. pneumonia and healthy vs. COVID-19; p < 0.001 for all; **Fig. 6-B**), whereas nasal swab and *’Unknown’* contributions remained unchanged. These findings align with our MaAsLin3 results, where disease-associated taxa losses were primarily prevalence-driven, and collectively support a model of disrupted microbial continuity and impaired dispersal from upper compartments during infection.

**Figure 6.**
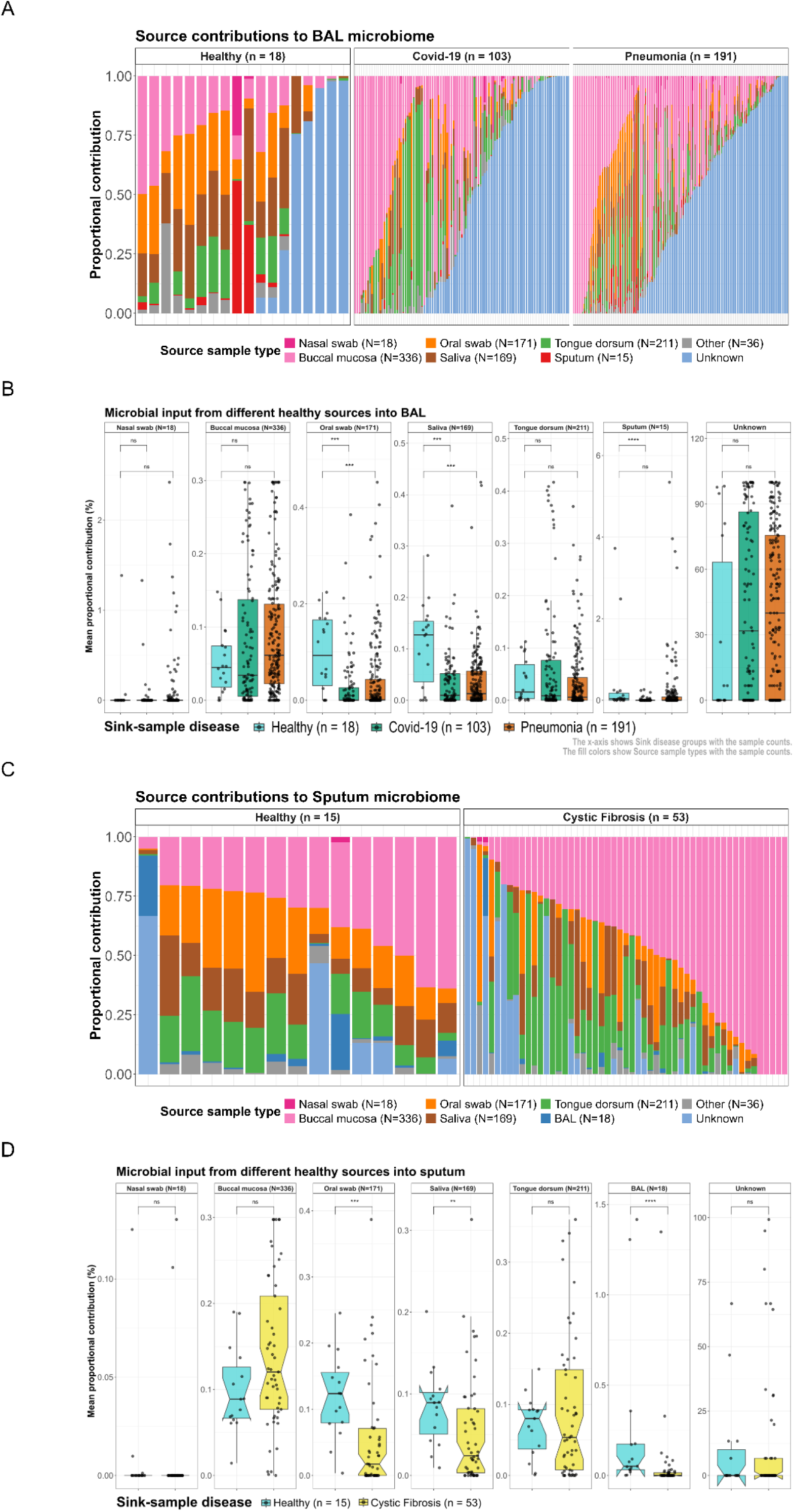
Microbial source contributions to the lower and intermediate respiratory tract in health and disease. (A) Proportional source contributions to BAL samples—used as proxies for the lower respiratory tract (LRT)—in healthy individuals (n = 19), and patients with pneumonia (n = 194) or COVID-19 (n = 103). Healthy BAL samples received input from a range of URT and IRT sites (notably tongue dorsum, oral swabs, and buccal mucosa), while disease samples showed a consistent reduction in known source contributions and a relative enrichment of unassigned (unknown) sources. (B) Boxplots displaying the distribution of source contributions to BAL across health and disease states. Source inputs from most URT and IRT sites were significantly reduced in pneumonia and COVID-19 compared to healthy samples (*P* < 0.001, Wilcoxon rank-sum), with nasal swab and “Unknown” sources showing no significant difference across groups. (C) Proportional source contributions to sputum samples (used as proxies for the intermediate respiratory tract, IRT) inferred from microbial source tracking in healthy individuals (n = 15) and patients with CF (n = 53). In CF, sputum communities were dominated by taxa originating from oral swabs and buccal mucosa, while healthy individuals showed more diverse and balanced inputs from multiple upper respiratory tract (URT) sources. (D) Boxplots showing source-specific contribution distributions to sputum across groups. Notable decreases in oral swab and saliva sources were observed in CF compared to healthy individuals (Wilcoxon rank-sum, *P* < 0.001 and *P* < 0.01, respectively).

We next applied source tracking to sputum samples from healthy and cystic fibrosis (CF) individuals to explore microbial dispersal into the IRT. While healthy sputum microbiota showed heterogeneous input from multiple upper airway sites, CF samples exhibited more individualized and irregular source profiles, often marked by an elevated proportion of oral microbial origins (**Fig. 6-C**). Contributions from oral swab samples and saliva were significantly reduced in CF sputum (q < 0.0001 for both, respectively; **Fig. 6-D**), challenging the expectation of increased oral influx in disease. These patterns suggest that CF-associated dysbiosis in the IRT may reflect not just altered input, but also selective filtering or survival favoring non-oral or uncharacterized taxa.

Together, these results reveal that respiratory diseases such as pneumonia, COVID-19, and CF are associated with a disruption in the spatial continuity of the respiratory microbiome, marked by reduced contributions from canonical source niches and a rise in microbial taxa of unknown origin. This breakdown in source–sink relationships highlights disease-associated impairments in microbial dispersal, colonization, or survival – offering ecological context to the reproducible prevalence losses observed across taxa in disease states.

## Discussion

In this study, we assembled and uniformly processed a large-scale, multi-cohort collection of human respiratory tract (RT) metagenomes to investigate microbial shifts across anatomical sites and disease states. By stratifying samples and applying batch-aware multivariable modeling, we disentangled patterns linked to the lower (LRT) and intermediate (IRT) respiratory tract in pneumonia, COVID-19, and cystic fibrosis (CF). The resulting open-access dataset provides a foundational resource for systems-level investigations of respiratory microbial dynamics.

Anterior nares samples formed a strikingly distinct cluster, dominated by high-abundance taxa such as *Staphylococcus* and *Corynebacterium* - consistent with prior reports from the Human Microbiome Project and other surveys^22,26–28^. Their compositional simplicity and individuality underscore the nares as a unique ecological niche, justifying their exclusion from broader RT analyses where they would obscure subtle gradients across the upper and lower tract.

A central finding is that respiratory disease is characterized less by shifts in relative abundance than by widespread losses in species prevalence. In pneumonia BAL samples, over 100 taxa showed reduced prevalence, reflecting collapse of community structure and loss of core commensals. Similar prevalence-driven disruption was seen in cystic fibrosis sputum, where only rare enrichments such as *Staphylococcus aureus* emerged; notably, *S. aureus* is a persistent colonizer of the nasal microbiota, often dominating specific community state types alongside *S. epidermidis*^28^. Notably, oral-associated taxa such as *Rothia mucilaginosa* and *Prevotella melaninogenica*, typically regarded as core species in healthy oral and airway niches, were consistently depleted across disease cohorts^29,30^.

These patterns align with a “loss of commensals” model, where taxa become less common across individuals and less abundant overall, rather than uniform pathogen expansion^31^. Although sporadic dominance events were observed, they were rare and lacked statistical significance. Prevalence-driven losses likely reflect combined pressures of host immunity, antibiotics, and impaired mucociliary clearance, which constrain colonization stability^32–34^. Crucially, these shifts were often invisible in abundance-only analyses, highlighting the value of prevalence modeling for capturing disease-associated dysbiosis in low-biomass ecosystems such as the lung^1,4^.

Source tracking analysis confirmed the view that, in health, the URT serves as a key microbial reservoir for the LRT. In these cases, the lung microbiome, represented by BAL samples, receives diverse and traceable contributions from URT and IRT sites, e.g., by microaspiration^35,36,1^, particularly the oral cavity, buccal mucosa, and tongue dorsum. This pattern is consistent with the concept that the healthy lung hosts a transient and spatially structured microbial community, maintained by continuous seeding through microaspiration or mucosal dispersion, and tightly regulated by host defense mechanisms such as mucociliary clearance, coughing, and immune surveillance^37^.

### Disruption of microbial continuity in disease

In pneumonia and COVID-19, this continuity collapsed: URT contributions to the LRT diminished, while unassigned microbial origins increased. Rather than being seeded by canonical URT reservoirs, the LRT microbiome in disease reflects selective colonization by more resilient or opportunistic taxa, or contributions from alternative and currently uncharacterized microbial sources. These changes likely result from disease-driven disruptions in host barrier function, immune responses, or ecological filtering in the inflamed lung environment^3,8,13^.

Our findings echo and extend previous reports suggesting that the healthy lower respiratory tract does not harbor a completely distinct microbiota but instead reflects a subset of upper airway communities, consistent with a topographical continuum, elsewhere also referred to as the adapted island model of lung biogeography^1,35^. This model sees the healthy RT as a single ecosystem, characterized by the constant dispersal and dilution of microbes from the URT to the LRT^35^. However, the observed taxa were not uniformly distributed, and disease states introduced distinct microbial shifts, indicating that selective ecological filtering, rather than passive dispersion alone, likely governs microbial colonization along the respiratory tract. These patterns challenge the notion of an undifferentiated lung microbiome and underscore the need for nuanced ecological models of respiratory microbial dynamics.

Despite the scale and integrative analysis of our analysis, several limitations should be considered and discussed. Despite rigorous batch correction and cohort-aware modeling, some taxa remained associated with specific bioprojects, indicating residual confounding by study design, geography, or technical factors. This highlights the need for future studies to adopt standardized sampling and comprehensive metadata reporting to enhance reproducibility in the future^38,39^. Disease-associated prevalence loss likely reflects cumulative effects of chronic inflammation, antibiotic exposure, and impaired mucus clearance^12,40,41^, but unmeasured host or environmental variables may also contribute. Multi-OMICs approaches with longitudinal sampling of clinically well-characterized cohorts could help to disentangle the various effects. Technical challenges remain acute in the LRT, where low microbial biomass increases susceptibility to contamination. BAL samples may capture upper-airway contaminants via bronchoscopy, while sputum inevitably mixes upper and lower tract material (REF). These caveats highlight the need for careful interpretation of anatomical specificity, particularly when comparing across disease cohorts and sampling strategies.

In summary, our study provides a novel foundational research for the respiratory tract research community, allowing for a comprehensive, multi-compartmental view of the human respiratory microbiome in health and disease. By integrating large-scale metagenomic data across anatomical sites and disease states, we reveal consistent patterns of microbial disruption, loss of commensals, and altered microbial connectivity in the diseased respiratory tract. These findings challenge static views of respiratory microbial ecology and underscore the need for dynamic, spatially resolved models to guide future diagnostics and therapeutic interventions.

## Methods

Overall, this study analyzed respiratory tract metagenomes from diverse populations, disease conditions, and anatomical sites. The workflow included standardized data retrieval, metadata harmonization, quality control, taxonomic profiling using MetaPhlAn4, and statistical modeling with batch-aware normalization to enable robust cross-cohort comparisons^42^.

### Metagenomic dataset collection

Metagenomic datasets were obtained from two primary sources: publicly available data from the NCBI Sequence Read Archive (SRA) and a contributed dataset generated for this study.

1 . NCBI-SRA open source metagenome data collection.

We queried the NCBI-SRA for publicly available human whole metagenome sequencing datasets originating from different regions of the respiratory tract using custom search strategies. We downloaded SRA run tables for each query with the basic sequencing info available on SRA. Selection of samples based on Library Layout (paired), sequencing technology (Illumina NGS), library preparation (random selection) with custom script in R (v.4.1.2)^43^, based on the following criteria: paired-end library layout, Illumina sequencing technology, and randomly selected library preparation protocols. Read counts were estimated using the ‘Bases’ variable and the average read length (‘AvgSpotLen’) to exclude samples with fewer than 100,000 reads. Redundant samples matching multiple queries were removed. We then compiled a list of samples with annotated sample types per BioProject to facilitate batch-wise download of read sequences. All samples meeting the selection criteria were subsequently downloaded using the SRA Toolkit (v3.0.0)^44^.

2. Contributed human metagenomic dataset

Additionally, a cohort of bronchoalveolar lavage (BAL) samples (N = 330) was contributed by co-author Tomas Eagan, comprising individuals with COPD, Asthma, Sarcoidosis, idiopathic pulmonary fibrosis, and healthy controls. These samples were collected as part of the MicroCOPD^45^ and MicroILD^20^ studies. Informed consent was obtained from all participants, and the study protocol was approved by the Helse-Vest and Helse-Nord regional ethical committees (IRB approval no. 2011/1307 (MicroCOPD) & 2014/1393 (MicroILD)). Sample collection and processing followed the protocols described in [reference or publication]. Samples were processed according to the protocol published in this paper^45,20,19^. All raw metagenomic reads sequenced for this cohort have been deposited in the European Nucleotide Archive (ENA) and are available under the study accession PRJEB96845

### Preprocessing and quality control of reads

To minimize taxonomic misclassification and spurious signals, we applied a strict multi-step quality control and preprocessing pipeline to the raw paired-end metagenomic reads, ensuring high-quality, host-DNA-decontaminated data for downstream analysis.

Human host-derived reads were removed by mapping the raw reads to the GRCh38 reference genome using *removehuman.sh* from the *BBMap* suite (v.39.01)^46^ with default parameters. Read quality filtering, adapter trimming, base correction, and complexity filtering were performed simultaneously using *fastp* (v0.23.4)^47^ with the following parameters: −− 𝑎𝑣𝑒𝑟𝑎𝑔𝑒_𝑞𝑢𝑎𝑙 20, −− 𝑑𝑒𝑡𝑒𝑐𝑡_𝑎𝑑𝑎𝑝𝑡𝑒𝑟_𝑓𝑜𝑟_𝑝𝑒, −− 𝑐𝑜𝑟𝑟𝑒𝑐𝑡𝑖𝑜𝑛, −− 𝑜𝑣𝑒𝑟𝑟𝑒𝑝𝑟𝑒𝑠𝑒𝑛𝑡𝑎𝑡𝑖𝑜𝑛_𝑎𝑛𝑎𝑙𝑦𝑠𝑖𝑠, and −− 𝑙𝑜𝑤_𝑐𝑜𝑚𝑝𝑙𝑒𝑥𝑖𝑡𝑦_𝑓𝑖𝑙𝑡𝑒𝑟.

To account for any unpaired reads resulting from trimming or filtering steps, we applied BBMap’s *repair.sh* script to restore proper read pairing^46^. Parameters included 𝑖𝑔𝑛𝑜𝑟𝑒𝑏𝑎𝑑𝑞𝑢𝑎𝑙𝑖𝑡𝑦 = 𝑡, 𝑡𝑜𝑠𝑠𝑏𝑟𝑜𝑘𝑒𝑛𝑟𝑒𝑎𝑑𝑠 = 𝑡, and 𝑜𝑣𝑒𝑟𝑤𝑟𝑖𝑡𝑒 = 𝑡. This ensured compatibility with downstream tools that require synchronized read pairs.

All quality control steps were performed using open-source software and parallelized across different HPC clusters with appropriate numbers of computational threads to improve efficiency. Quality assessments were performed both before and after preprocessing using *MultiQC*^48^ aggregated reports from *FastQC* (v0.11.9)^49^, evaluating metrics such as per-base quality scores and adapter contamination. Samples with fewer than 100,000 reads after quality control were excluded from downstream analysis.

### Metadata curation and harmonization

Sample-associated metadata were manually curated from original publications, NCBI BioSample records, and supplementary tables. Considerable effort was taken to standardize and harmonize variable formats across datasets. Only variables consistently available across all cohorts were retained for downstream analysis. Additionally, a composite variable *RT_category* was created to classify samples based on respiratory tract location, which was appended to the unified metadata table. Although not anatomically defined as a discrete compartment, within the *RT_category* variable, we use the term ‘intermediate respiratory tract’ (IRT) to group sampling sites that bridge the upper and lower respiratory tract (such as palatine tonsils, tongue dorsum, supraglottic, throat, and sputum), consistent with their transitional anatomical position and characteristic microbial profiles. Definitions of all metadata variables are provided in Supplementary Table 2.

#### Metagenomic taxonomic profiling and microbiome analysis

Taxonomic profiling was performed using MetaPhlAn (v4.1.1)^42^, implemented via the AnADAMA2^50^ workflow framework, leveraging the latest marker gene database (*vOct22_202403*) to estimate species-level microbial composition from metagenomic reads. MetaPhlAn profiling identified 261 samples with entirely unclassified taxonomic profiles; these were excluded from downstream analyses.

Alpha and beta diversity metrics were computed using a custom R script. Alpha diversity indices - including observed richness, Shannon, Simpson, and Gini dominance - were calculated using the microbiome R package (v.1.26)^51^. Beta diversity was estimated using Bray-Curtis^52^, Jaccard^53^, weighted and unweighted UniFrac^54^, and Aitchison^55^ distances via the rbiom (v1.0.3)^56^, compositions (v2.0.8)^57^, and ape (v5.8.1)^58^ packages. Bray-Curtis dissimilarity matrices were visualized using Principal Coordinates Analysis (PCoA). Variance explained by the top components was used to interpret community structure, and outliers were identified based on Z-scores (|z| > 2.5) calculated from the first two principal coordinates. Samples flagged as outliers were excluded from downstream analysis to avoid distortion in diversity and association results.

To assess the individual contribution of sample metadata variables to microbial community composition, univariate PERMANOVA^59^ tests were performed using the *adonis2()* function from the vegan R package (v2.6.8)^60^. Bray–Curtis dissimilarity matrices calculated from species-level taxonomic profiles were used as input. R² values and permutation-based p-values (n = 10,000) were extracted for each model to quantify effect size and statistical significance.

#### Batch correction across studies

Taxa relative abundance profiles (from MetaPhlAn) were batch-corrected using the adjust_batch() function from the MMUPHin (v1.20)^61^ R package to account for study-specific batch effects. Correction was performed within each RT category by subsetting samples accordingly. For the IRT, only sputum samples were included, as it was the only sample type shared across multiple BioProjects. Taxa present in fewer than two samples were excluded before correction to reduce sparsity and improve model stability.

#### Differential prevalence/abundance analysis in the lower and intermediate respiratory tract

To identify microbial taxa associated with respiratory diseases, we applied MaAsLin 3 (v3.0.0)^62^ for multivariable modeling of microbial features, using species-level, batch-corrected relative abundance profiles derived from MetaPhlAn 4^42^. Analyses were performed separately for BAL and sputum samples, enabling compartment-specific modeling of the lower and intermediate respiratory tract. Only healthy and disease-specific subsets were used to minimize confounding by sample type.

Species-level feature tables were filtered to retain taxa detected in at least two samples, and samples with extremely low observed alpha diversity (observed diversity <2; two or fewer species) were excluded to avoid artifacts from low-biomass or contaminated samples. Before modeling, microbial profiles were normalized using total sum scaling (TSS) and log-transformed. The primary model tested associations between disease status (*Healthy*, *Pneumonia*, *COVID* − 19, or *Cystic Fibrosis*) and microbial features, while controlling for sequencing depth (*Total_reads*): Cross-cohort Model 1: ∼ 𝐷𝑖𝑠𝑒𝑎𝑠𝑒 + *Total_reads*

This was applied separately to:

● BAL samples for comparisons across *Healthy*, *Pneumonia*, and *COVID* − 19 individuals.
● Sputum samples for comparisons between *Healthy* and *Cystic Fibrosis* individuals.

To assess the robustness and reproducibility of species–disease associations, we complemented this with within-dataset models (Model 2), where the same formula was applied separately to each disease cohort. This cohort-aware approach allowed us to identify species consistently associated with disease across independent datasets, and to flag potential cohort-specific signals. Species that showed significant associations with *Total_reads* were flagged as potentially confounded and treated with caution in interpretation.

All models were run using 6 CPU cores and the following parameters:𝑛𝑜𝑟𝑚𝑎𝑙𝑖𝑧𝑎𝑡𝑖𝑜𝑛 = ’𝑇𝑆𝑆’, 𝑡𝑟𝑎𝑛𝑠𝑓𝑜𝑟𝑚 = ’𝐿𝑂𝐺’, and 𝑠𝑡𝑎𝑛𝑑𝑎𝑟𝑑𝑖𝑧𝑒 = 𝑇𝑅𝑈𝐸. Species with FDR-adjusted q-values < 0.05 were considered statistically significant and further ranked by their effect size (*β*-coefficient) and sample prevalence. To enable direct visual comparison of global vs. cohort-specific effects, taxa with consistent associations in both the main (Model 1) and auxiliary models (Model 2) were annotated on coefficient plots using heatmap-style overlays.

#### Microbial source tracking across the respiratory tract

To investigate the directional flow and potential microbial contributions between anatomical sites within the respiratory tract, source tracking analysis was performed using the FEAST tool (v3.6.5)^63^. Analyses were conducted separately for healthy and diseased individuals. For each group, taxa present in at least 10 samples were retained, and relative abundances were averaged across samples from the same individual to reduce intra-host noise. FEAST was then used to estimate the proportional contribution of URT and IRT communities to LRT samples. Contributions were aggregated by sample type (e.g., saliva, sputum, BAL) to assess source-specific influence via group-wise comparisons and visualized using boxplots.

#### Statistics and visualization

All data cleaning, statistical analysis, and visualization were performed in R (v4.3) unless otherwise specified. Statistical modeling, including multivariable regression of microbial features, was conducted using the R package MaAsLin 3 (v3.0.0) with custom R scripts for feature filtering, significance evaluations, and plotting results. Beta diversity metrics (Bray–Curtis, Jaccard, UniFrac, and Aitchison) were computed using the rbiom (v1.0.3), microbiome (v1.26), ape (v5.8.1), and compositions (v2.0.8) packages. Principal Coordinates Analysis (PCoA) and PERMANOVA (adonis2) were carried out using vegan (v2.6-8). Significance stars, axis ordering, and taxon filtering were added through custom plotting logic to enhance interpretability.

## Data availability

The raw sequencing reads generated and analyzed in this study have been deposited in the European Nucleotide Archive (ENA) under project accession PRJEB96845. In addition, we re-analysed previously published datasets available in the ENA/NCBI Sequence Read Archive under the following BioProject accessions: PRJNA48479 and PRJNA917836. A complete list of all reused BioProjects with sample counts and sequencing statistics is provided in **Table S-1**. All custom code and analysis workflows are available in a GitHub repository (https://github.com/CME-lab-research/MiPoRT). Processed taxonomic profiles, supplementary tables and file, and are also available here.

## Acknowledgements

The following resources are gratefully acknowledged. The computational analysis in this work were run on different platforms: MedBioNode run by the Core Facility Computational Bioanalytics (Medical University of Graz, funded by the Austrian Federal Ministry of Education, Science and Research, Hochschulraum-Strukturmittel 2016 grant as part of BioTechMed Graz), LiSC (University of Vienna), FASRC Cannon cluster, as provided by the FAS Division of Science Research Computing Group (Harvard University, Boston, MA). This work was financially supported by the Medical University of Graz and the Austrian Science Fund (FWF) through the following programs: doc. funds RESPImmun (Grant DOI 10.55776/DOC129, given to CME (Co-PI) and Grazyna Kwapiszewska (PI)), Cluster of Excellence “Microbiomes drive Planetary Health” (Grant DOI 10.55776/COE7, given to CME (Co-PI)), and SFB “Immunometabolism” (Grant DOI 10.55776/F8300, given to CME (Co-PI)). We further acknowledge the financial support by MEFO Graz. The research stay of TS in the CH lab was funded by travel grants obtained from the Austrian Marshall Plan foundation and the Medical University of Graz “Microbiome & Infection” Research Field. VF is supported through a Mid-Career Grant of the Austrian Society of Pneumology and through a Max-Kade Fellowship. We are grateful for the scientific support (discussions on study layout and methodology) provided by Nicola Segata and Stefanie Widder, who served together with VF as thesis committee members for TS. We thank Klara Filek, and Christian Diener for valuable discussions and feedback, and acknowledge the RESPImmun PhD program for providing a stimulating training environment.

## Authors’ contributions

TS and CME conceived the study together with VF and CK. TS performed the formal analyses and, with CME, VF, TE, YZ, EF, and CH, developed the methodology. CME and CK coordinated project administration. Supervision was provided by CME, VF, CK, CH, YZ, and EF. TE provided original data and, along with VF, clinical expertise. RM and VW contributed to critical discussions and figures. TS drafted the original manuscript, and all authors contributed to the review and editing of the final version.

## References

1. Dickson, R. P., Erb-Downward, J. R., Martinez, F. J. & Huffnagle, G. B. The Microbiome and the Respiratory Tract. Annu. Rev. Physiol. 78, 481–504 (2016).

2. Hewlett, R. T. THE FATE OF MICRO-ORGANISMS IN INSPIRED AIR. The Lancet 147, 86–87 (1896).

3. Huffnagle, G. B. & Dickson, R. P. The bacterial microbiota in inflammatory lung diseases. Clin. Immunol. 159, 177–182 (2015).

4. Man, W. H., de Steenhuijsen Piters, W. A. A. & Bogaert, D. The microbiota of the respiratory tract: gatekeeper to respiratory health. Nat. Rev. Microbiol. 15, 259–270 (2017).

5. Pérez-Cobas, A. E., Rodríguez-Beltrán, J., Baquero, F. & Coque, T. M. Ecology of the respiratory tract microbiome. Trends Microbiol. 31, 972–984 (2023).

6. Charlson, E. S. et al. Topographical Continuity of Bacterial Populations in the Healthy Human Respiratory Tract. Am. J. Respir. Crit. Care Med. 184, 957–963 (2011).

7. Durack, J. et al. Bacterial biogeography of adult airways in atopic asthma. Microbiome 6, 104 (2018).

8. Mika, M. et al. Microbial and host immune factors as drivers of COPD. ERJ Open Res. 4, (2018).

9. Hilty, M. et al. Disordered Microbial Communities in Asthmatic Airways. PLOS ONE 5, e8578 (2010).

10. Einarsson, G. G. et al. Community dynamics and the lower airway microbiota in stable chronic obstructive pulmonary disease, smokers and healthy non-smokers. (2016) doi:10.1136/thoraxjnl-2015-207235.

11. Rick, E.-M. et al. The airway fungal microbiome in asthma. Clin. Exp. Allergy 50, 1325–1341 (2020).

12. Belkaid, Y. & Hand, T. W. Role of the Microbiota in Immunity and Inflammation. Cell 157, 121–141 (2014).

13. Gilbert, J. A. et al. Current understanding of the human microbiome. Nat. Med. 24, 392–400 (2018).

14. Kumpitsch, C., Koskinen, K., Schöpf, V. & Moissl-Eichinger, C. The microbiome of the upper respiratory tract in health and disease. BMC Biol. 17, 87 (2019).

15. Loverdos, K. et al. Lung Microbiome in Asthma: Current Perspectives. J. Clin. Med. 8, 1967 (2019).

16. Luo, L., Tang, J., Du, X. & Li, N. Chronic obstructive pulmonary disease and the airway microbiome: A review for clinicians. Respir. Med. 225, 107586 (2024).

17. Cuthbertson, L. et al. Lung function and microbiota diversity in cystic fibrosis. Microbiome 8, 45 (2020).

18. Widder, S. et al. Microbial community organization designates distinct pulmonary exacerbation types and predicts treatment outcome in cystic fibrosis. Nat. Commun. 15, 4889 (2024).

19. Knudsen, K. S. et al. The lower airways microbiota and antimicrobial peptides indicate dysbiosis in sarcoidosis. Microbiome 10, 175 (2022).

20. Knudsen, K. S. et al. The lower airways microbiome and antimicrobial peptides in idiopathic pulmonary fibrosis differ from chronic obstructive pulmonary disease. PLOS ONE 17, e0262082 (2022).

21. Zhu, T., Jin, J., Chen, M. & Chen, Y. The impact of infection with COVID-19 on the respiratory microbiome: A narrative review. Virulence 13, 1076–1087 (2022).

22. Huttenhower, C. et al. Structure, function and diversity of the healthy human microbiome. Nature 486, 207–214 (2012).

23. Horz, H. P., Robertz, N., Vianna, M. E., Henne, K. & Conrads, G. Relationship between methanogenic archaea and subgingival microbial complexes in human periodontitis. Anaerobe 35, 10–12 (2015).

24. Vashishta, A. et al. Filifactor alocis Pathogenicity Requires TLR2 and the Oral Microbiome. J. Dent. Res. 00220345251331959 (2025) doi:10.1177/00220345251331959.

25. Laguna, T. A. et al. Airway Microbiota in Bronchoalveolar Lavage Fluid from Clinically Well Infants with Cystic Fibrosis. PLOS ONE 11, e0167649 (2016).

26. De Boeck, I., et al. Anterior Nares Diversity and Pathobionts Represent Sinus Microbiome in Chronic Rhinosinusitis. mSphere 4, 10.1128/msphere.00532-19 (2019).

27. Cole, A. L. et al. Identification of Nasal Gammaproteobacteria with Potent Activity against Staphylococcus aureus: Novel Insights into the “Noncarrier” State. mSphere 6, e01015–20 (2021).

28. Ingham, A. C. et al. Staphylococci in high resolution: Capturing diversity within the human nasal microbiota. Cell Rep. 44, 115854 (2025).

29. Uranga, C. C., Arroyo, P., Duggan, B. M., Gerwick, W. H. & Edlund, A. Commensal Oral Rothia mucilaginosa Produces Enterobactin, a Metal-Chelating Siderophore. mSystems 5, 10.1128/msystems.00161-20 (2020).

30. Könönen, E., Fteita, D., Gursoy, U. K. & Gursoy, M. Prevotella species as oral residents and infectious agents with potential impact on systemic conditions. J. Oral Microbiol. 14, 2079814.

31. Neville, B. A., Forster, S. C. & Lawley, T. D. Commensal Koch’s postulates: establishing causation in human microbiota research. Curr. Opin. Microbiol. 42, 47–52 (2018).

32. Surette, M. G. The Cystic Fibrosis Lung Microbiome. Ann. Am. Thorac. Soc. 11, S61–S65 (2014).

33. Knoll, R. L. et al. Resilience and stability of the CF- intestinal and respiratory microbiome during nutritional and exercise intervention. BMC Microbiol. 23, 44 (2023).

34. Cauwenberghs, E. et al. Positioning the preventive potential of microbiome treatments for cystic fibrosis in the context of current therapies. Cell Rep. Med. 5, 101371 (2024).

35. Venkataraman, A. et al. Application of a Neutral Community Model To Assess Structuring of the Human Lung Microbiome. mBio 6, 10.1128/mbio.02284-14 (2015).

36. Dickson, R. P. et al. Spatial Variation in the Healthy Human Lung Microbiome and the Adapted Island Model of Lung Biogeography. Ann. Am. Thorac. Soc. 12, 821–830 (2015).

37. Aho, V. T. E. et al. The microbiome of the human lower airways: a next generation sequencing perspective. World Allergy Organ. J. 8, 23 (2015).

38. Sinha, R., Abnet, C. C., White, O., Knight, R. & Huttenhower, C. The microbiome quality control project: baseline study design and future directions. Genome Biol. 16, 276 (2015).

39. Knight, R. et al. Best practices for analysing microbiomes. Nat. Rev. Microbiol. 16, 410–422 (2018).

40. 40. Global genetic diversity of human gut microbiome species is related to geographic location and host health: Cell. https://www.cell.com/cell/fulltext/S0092-8674(25)00416-7.

41. Stevens, J. et al. Microbiota-derived inosine programs protective CD8+ T cell responses against influenza in newborns. Cell (2025) doi:10.1016/j.cell.2025.05.013.

42. Blanco-Míguez, A. et al. Extending and improving metagenomic taxonomic profiling with uncharacterized species using MetaPhlAn 4. Nat. Biotechnol. 41, 1633–1644 (2023).

43. R Core Team. R: A Language and Environment for Statistical Computing. R Foundation for Statistical Computing (2021).

44. National Center for Biotechnology Information (NCBI). SRA Toolkit. (2022).

45. Grønseth, R. et al. The Bergen COPD microbiome study (MicroCOPD): rationale, design, and initial experiences. Eur. Clin. Respir. J. 1, 26196 (2014).

46. Bushnell, B., Rood, J. & Singer, E. BBMerge – Accurate paired shotgun read merging via overlap. PLOS ONE 12, e0185056 (2017).

47. Chen, S., Zhou, Y., Chen, Y. & Gu, J. fastp: an ultra-fast all-in-one FASTQ preprocessor. Bioinformatics 34, i884–i890 (2018).

48. Ewels, P., Magnusson, M., Lundin, S. & Käller, M. MultiQC: summarize analysis results for multiple tools and samples in a single report. Bioinformatics 32, 3047–3048 (2016).

49. Andrews, S. FastQC: A quality control tool for high throughput sequence data. Babraham Bioinformatics (2010).

50. McIver, L. J. AnADAMA2: Another Automated Data Analysis Management Application. (2015).

51. Lahti, L., Shetty, S. & Blake, T. microbiome: Tools for microbiome analysis in R. (2017).

52. Bray, J. R. & Curtis, J. T. An Ordination of the Upland Forest Communities of Southern Wisconsin. Ecol. Monogr. 27, 325–349 (1957).

53. Jaccard, P. Étude comparative de la distribution florale dans une portion des Alpes et du Jura. Bull. Société Vaudoise Sci. Nat. 37, 547 (1901).

54. Lozupone, C. & Knight, R. UniFrac: a New Phylogenetic Method for Comparing Microbial Communities. Appl. Environ. Microbiol. 71, 8228–8235 (2005).

55. Aitchison, J. The Statistical Analysis of Compositional Data. J. R. Stat. Soc. Ser. B Methodol. 44, 139–160 (1982).

56. Smith, D. P. rbiom: Read/Write, Analyze, and Visualize ‘BIOM’ Data. R package version v1.0.3, https://github.com/cmmr/rbiom. (2024).

57. van den Boogaart, K. G. & Tolosana-Delgado, R. compositions: Compositional Data Analysis.

58. Paradis, E. & Schliep, K. ape 5.0: an environment for modern phylogenetics and evolutionary analyses in R. Bioinformatics 35, 526–528 (2019).

59. Anderson, M. J. A new method for non-parametric multivariate analysis of variance. Austral Ecol. 26, 32–46 (2001).

60. Oksanen, J. et al. vegan: Community Ecology Package (v2.6.8). (2025).

61. Ma, S. et al. Population structure discovery in meta-analyzed microbial communities and inflammatory bowel disease using MMUPHin. Genome Biol. 23, 208 (2022).

62. Nickols, W. A. et al. MaAsLin 3: Refining and extending generalized multivariable linear models for meta-omic association discovery. 2024.12.13.628459 Preprint at 10.1101/2024.12.13.628459 (2024).

63. Shenhav, L. et al. FEAST: fast expectation-maximization for microbial source tracking. Nat. Methods 16, 627–632 (2019).

