## Supplementary Materials for "Pan-microbiome analysis along the human respiratory axis reveals an ecological continuum in health and collapse in disease"

---

[1] Diagnostic and Research Institute of Hygiene, Microbiology and Environmental Medicine, Medical University of Graz, 8010 Graz, Austria

[2] Division of Respiratory Medicine, Department of Internal Medicine, Lung Research Cluster, Medical University of Graz, 8010 Graz, Austria

[3] Department of Biostatistics, Harvard T.H. Chan School of Public Health, 02115 Boston, USA

[4] Shenzhen Branch, Guangdong Laboratory of Lingnan Modern Agriculture, Genome Analysis Laboratory of the Ministry of Agriculture and Rural Affairs, Agricultural Genomics Institute at Shenzhen, Chinese Academy of Agricultural Sciences, Shenzhen, China

[5] Department of Thoracic Medicine, Haukeland University Hospital, 5021 Bergen, Norway

[6] Channing Division of Network Medicine, Department of Medicine, Brigham and Women's Hospital, Harvard Medical School, 02115 Boston, USA

[7] Broad Institute of Harvard and MIT, Cambridge, Massachusetts (MA).

[8] Department of Immunology and Infectious Diseases, Harvard T. H. Chan School of Public Health, Boston, MA.

[9] Harvard Chan Microbiome in Public Health Center, Harvard T. H. Chan School of Public Health, Boston, MA.

[10] BioTechMed-Graz, 8010 Graz, Austria

| **ProjectID** | **Total raw read count** (before QC) | **Total read count after decontamination** (host DNA removal) | **Total read count** (after QC) | **Mean diff.** (raw v/s QC) | **Percent difference** (raw v/s QC) | **Mean #reads** (raw) | **Mean #reads** (QC) | **Sample count** |
|---|---|---|---|---|---|---|---|---|
| **PRJEB27079** | 546078658 | 982422 | 982422 | 136274059 | 24.96 | 136519665 | 245606 | 4 |
| **PRJEB29011** | 3398595496 | 887998636 | 784443934 | 12690057 | 0.37 | 16498037 | 3807981 | 206 |
| **PRJEB9034** | 101764756 | 46723198 | 87359156 | 800312 | 0.79 | 5653598 | 4853287 | 18 |
| **PRJNA316056** | 360120000 | 16524488 | 10361398 | 29146551 | 8.09 | 30010000 | 863450 | 12 |
| **PRJNA316588** | 1200566212 | 67260563 | 43451068 | 64284175 | 5.35 | 66698123 | 2413949 | 18 |
| **PRJNA380727** | 4152740588 | 1720911646 | 1647306194 | 42464990 | 1.02 | 70385434 | 27920444 | 59 |
| **PRJNA413615** | 4760697502 | 4760257632 | 4689698830 | 258177 | 0.01 | 17311628 | 17053451 | 275 |
| **PRJNA470402** | 4087364474 | 1602924374 | 1532871736 | 44042979 | 1.08 | 70471802 | 26428824 | 60 |
| **PRJNA48479** | 62764202771 | 29349325994 | 29349325994 | 29941646 | 0.05 | 56240326 | 26298680 | 1128 |
| **PRJNA494034** | 363543946 | 345825942 | 282872078 | 983804 | 0.27 | 4433463 | 3449660 | 82 |
| **PRJNA516442** | 198606752 | 176252460 | 189102154 | 158410 | 0.08 | 3310113 | 3151703 | 60 |
| **PRJNA516870** | 378653120 | 376011438 | 375667020 | 38284 | 0.01 | 4854528 | 4816244 | 78 |
| **PRJNA644285** | 254436806 | 21658943 | 21509366 | 19410620 | 7.63 | 21203068 | 1792448 | 12 |
| **PRJNA655567** | 269598330 | 87014877 | 166820664 | 1684880 | 0.62 | 4419645 | 2734765 | 61 |
| **PRJNA687506** | 41418482646 | 11566316700 | 11379804510 | 142363404 | 0.34 | 196296127 | 53932723 | 211 |
| **PRJNA687506 URT** | 15228989013 | 4247949276 | 4712522752 | 178245191 | 1.17 | 258118458 | 79873267 | 62 |
| **PRJNA756530** | 28609014 | 22295824 | 22281224 | 103735 | 0.36 | 469001 | 365266 | 61 |
| **PRJNA757846** | 2369832128 | 104055334 | 98620004 | 41294766 | 1.74 | 43087857 | 1793091 | 55 |
| **PRJNA757846 URT** | 2311879552 | 633152686 | 625602364 | 29073745 | 1.26 | 39859993 | 10786248 | 58 |
| **PRJNA762218** | 2154848364 | 741963662 | 725702868 | 62136761 | 2.88 | 93689060 | 31552299 | 23 |
| **PRJNA917836** | 108724502552 | 108719460384 | 107837960966 | 1164970 | 0 | 142870569 | 141705600 | 765 |
| **UniBergen** | 1137457244 | 1137457244 | 385178988 | 2279632 | 0.2 | 3446841 | 1167210 | 330 |
| **Total read count** (Bill.) | **256.21** | **166.63** | **164.97** | | | | | **N = 3638** |

**S-1 Table.**

Summary of sequencing read counts and quality control statistics across projects. This table reports the total and mean read counts at each stage of preprocessing, including raw data, host DNA decontamination, and final quality-controlled (QC) reads. Mean and percent differences between raw and QC read counts are provided to assess the extent of data retained. The number of samples processed per project is also indicated. This read counts table is sorted by BioProject-ID name in alphabetical order.

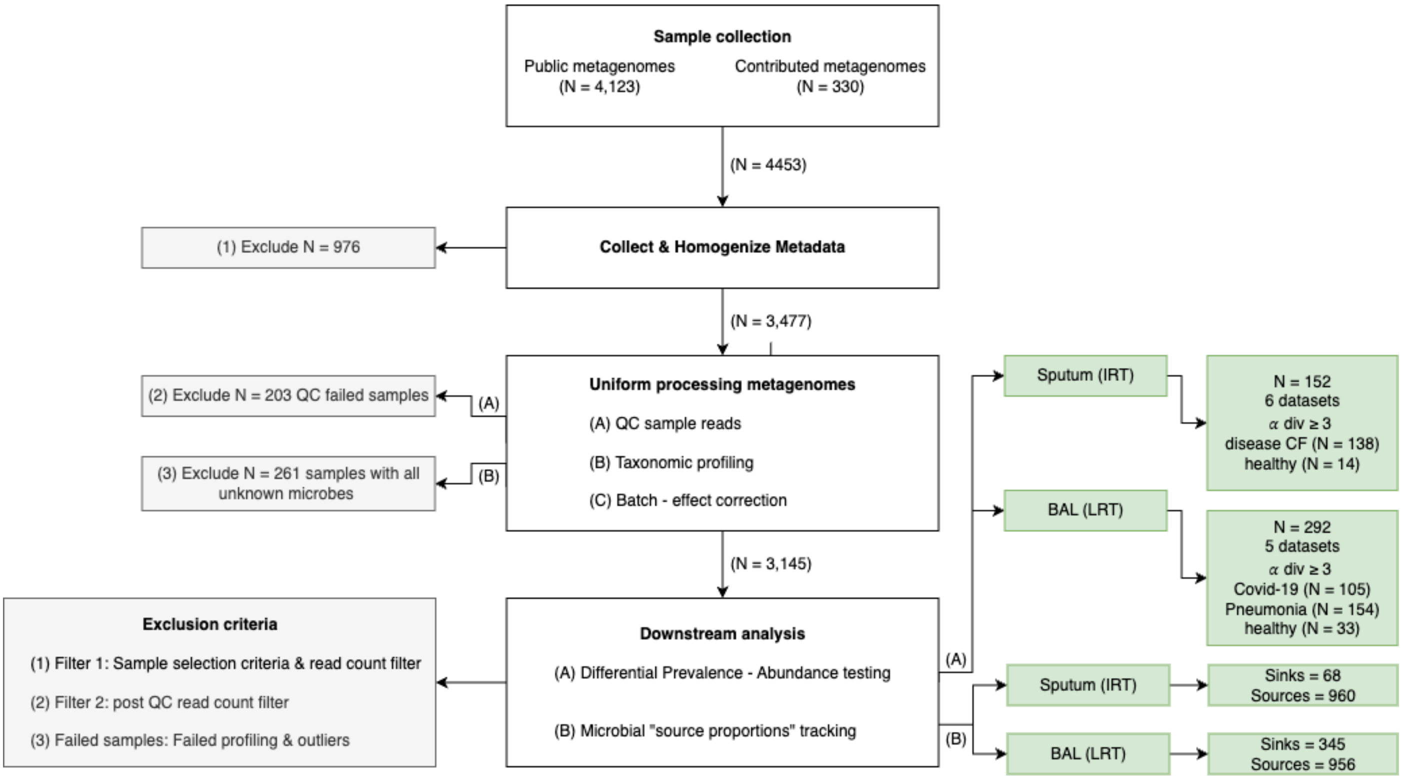


**S-1 Fig.**

Storm chart of data analysis steps and sample counts at each step (after filtering)

| Variable | Description | Type | Levels / Categories | Source | Privacy note |
|---|---|---|---|---|---|
| BioProjectID | Identifier for the sequencing project of origin | Categorical | e.g., PRJNA48479 | Public (NCBI-SRA) | Already public |
| SampleType | Anatomical sampling site | Categorical | Saliva, BAL, sputum, etc. | Study metadata | Non-sensitive |
| **RT_category *** | Grouped compartment (URT / IRT / LRT) | Categorical | URT, IRT, LRT | Harmonized & derived variable | Non-sensitive |
| Disease | Clinical diagnosis associated with the sample | Categorical | Healthy, Pneumonia, CF, COVID-19 | Study metadata | Non-sensitive (broad categories) |
| Healthy | Binary status for health | Categorical | TRUE, FALSE | Harmonized & derived variable | Non-sensitive (broad categories) |
| **AgeGroup **** | Age binned into categories | Ordinal | Child, Young-adult, Adult, Older-adult | Harmonized metadata variable | Aggregated for privacy |
| Sex | Reported biological sex | Categorical | Female, Male, NA | Harmonized metadata variable | Non-sensitive |
| Country | The country where the study was conducted | Categorical | e.g., Germany, Norway, China, USA, etc. | Public (NCBI-SRA) BioProject metadata | Already public |
| Patient-ID | Repeated measures subject identifier within the cohort | String | Anonymized codes (e.g., BioProjectID-S1, BioProjectID-S2) | Harmonized metadata variable | Anonymized within-study identifiers |
| Time_point | Sampling time for repeated samples from the same individual | Categorical | e.g., Time_pt-1, Time_pt-2, Visit_1, etc. | Study metadata | Non-sensitive (coded time points, not actual dates) |

**S-2 Table.**

Metadata variables harmonized across cohorts in the study, including their type, levels, source of origin, and privacy considerations. Derived composite variables ($RT\_category$, $AgeGroup$) were created to enable consistent cross-cohort comparisons. $RT\_category$ was aggregated as follows: **URT** (Upper RT) – anterior nares, nasal swab, nasopharyngeal aspirate, buccal mucosa, oral swab, saliva; **IRT** (Intermediate RT) – palatine tonsils, tongue dorsum, supraglottal, throat, sputum; **LRT** (Lower RT) – BAL, lung tissue. $AgeGroup$ were binned as Child (<16), Young-adult (16–30), Adult (30–45), Older-adult (>45), and NA (not available).

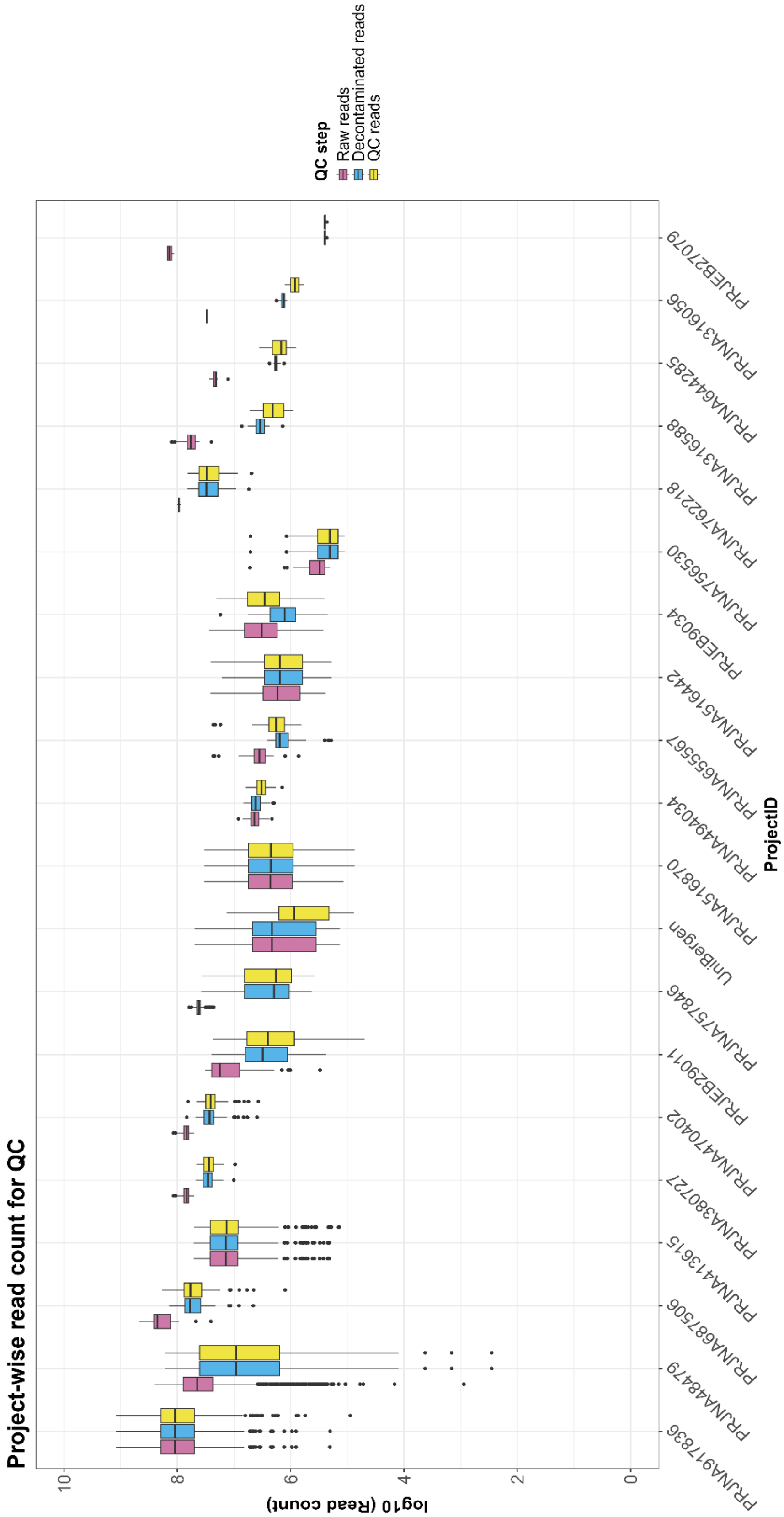


### S-2 Fig.

QC stepwise read count plot showing the number of reads retained for each Bio-project in this study.

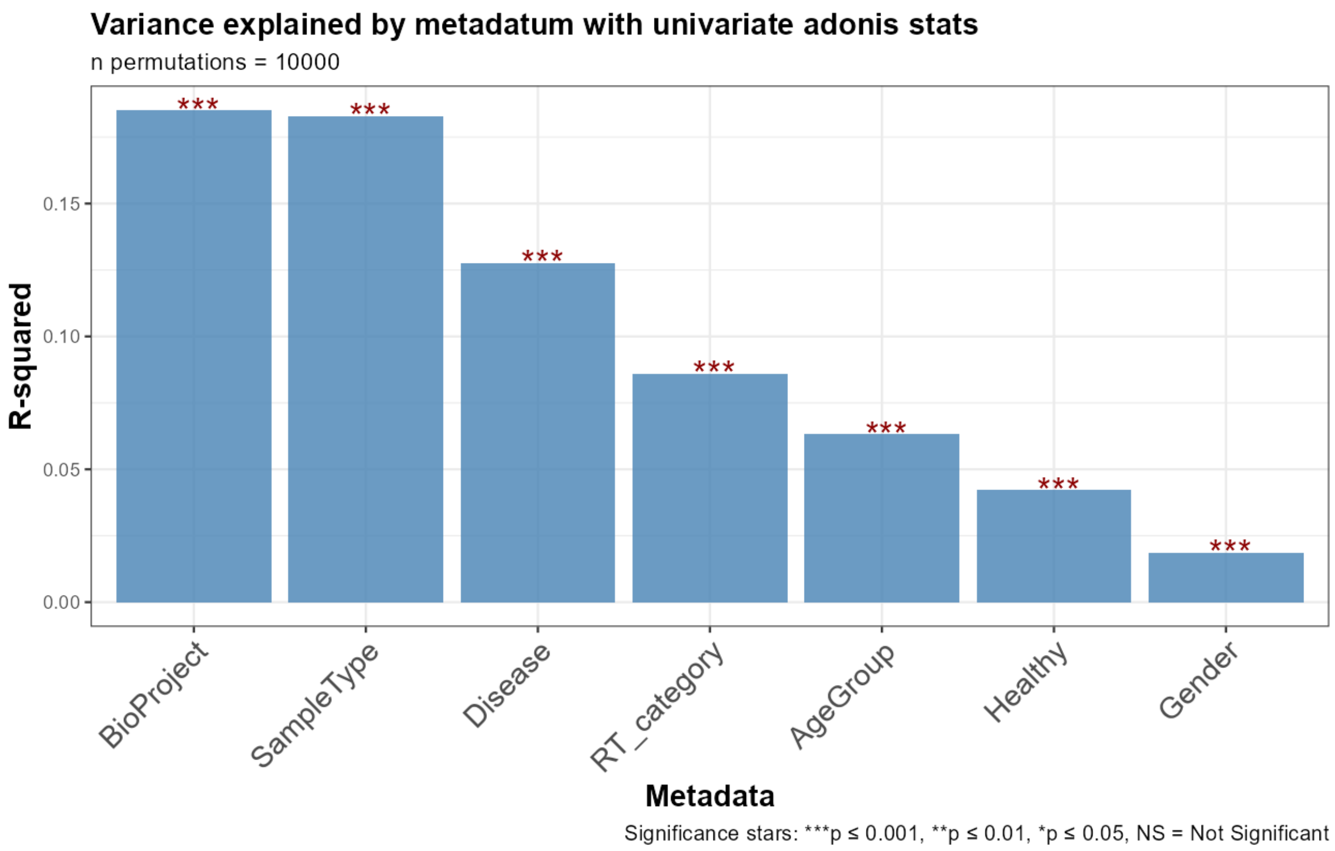


**S-3 Fig.**

Variance in microbial community structure explained by individual metadata variables before batch effect correction. Bar plot showing the proportion of variance ($R^2$) explained by each metadata variable in univariate PERMANOVA (adonis2) models based on Bray–Curtis dissimilarity. All p-values were calculated using permutation tests (10,000 permutations), unless otherwise specified.

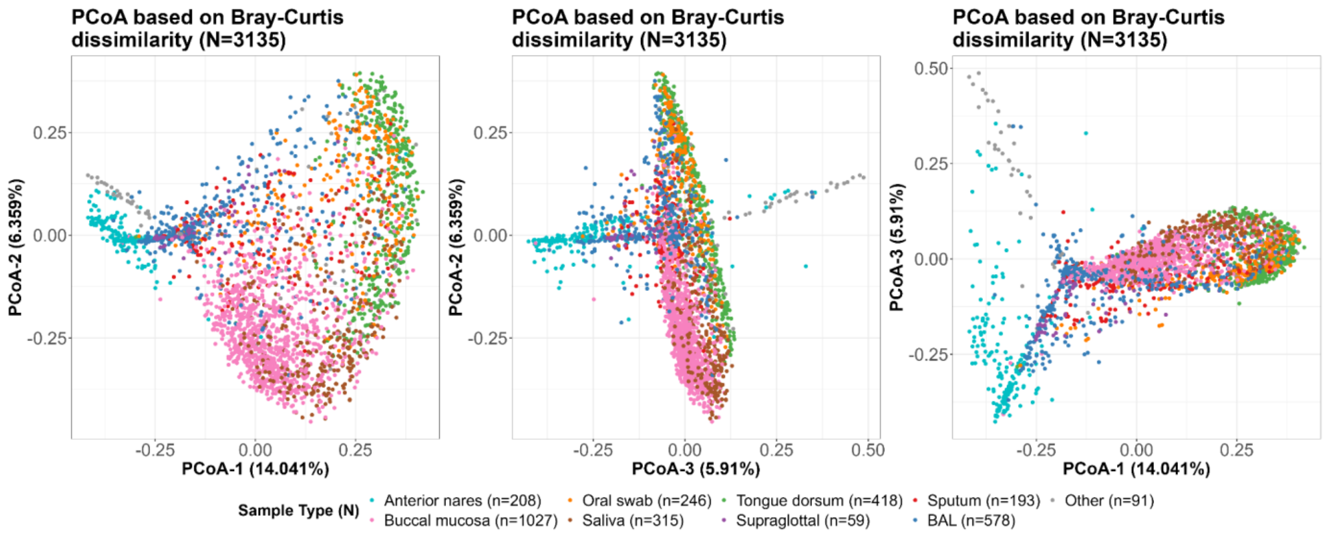


### S-4 Fig.

Ordination plot showing beta diversity (Bray-Curtis) of all analyzed samples, colored by sample type. The three plots display the top three principal components, which explain the maximum variation.

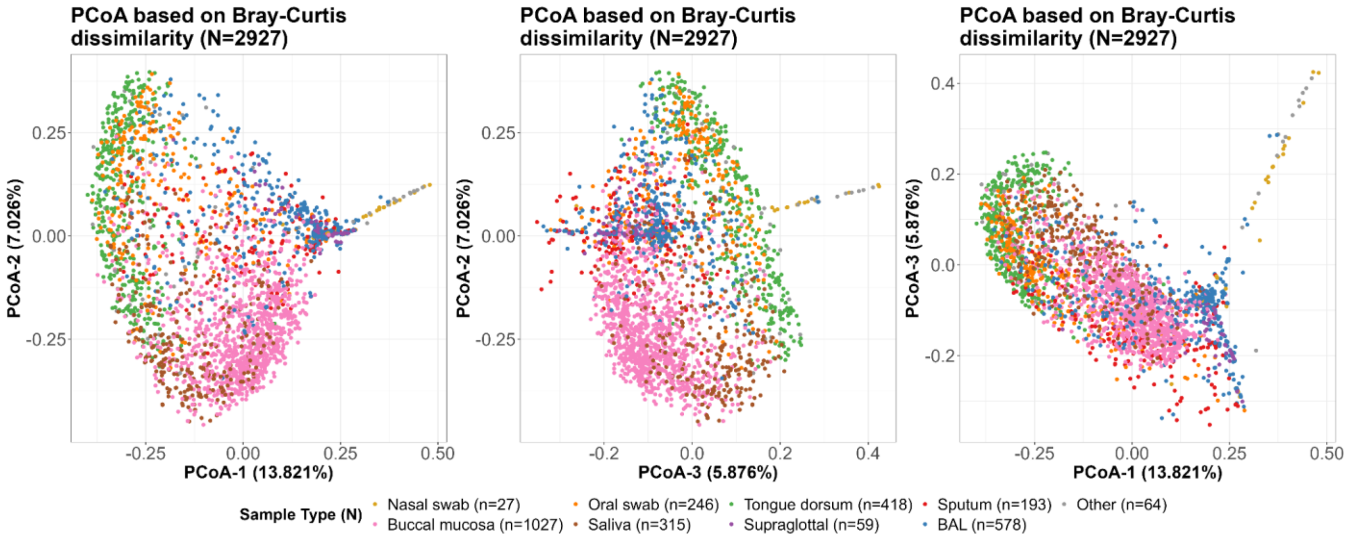


### S-5 Fig.

Ordination plot showing beta diversity (Bray-Curtis) of samples excluding Anterior nares, colored by sample type. The three plots display the top three principal components, which explain the maximum variation.

**A.**

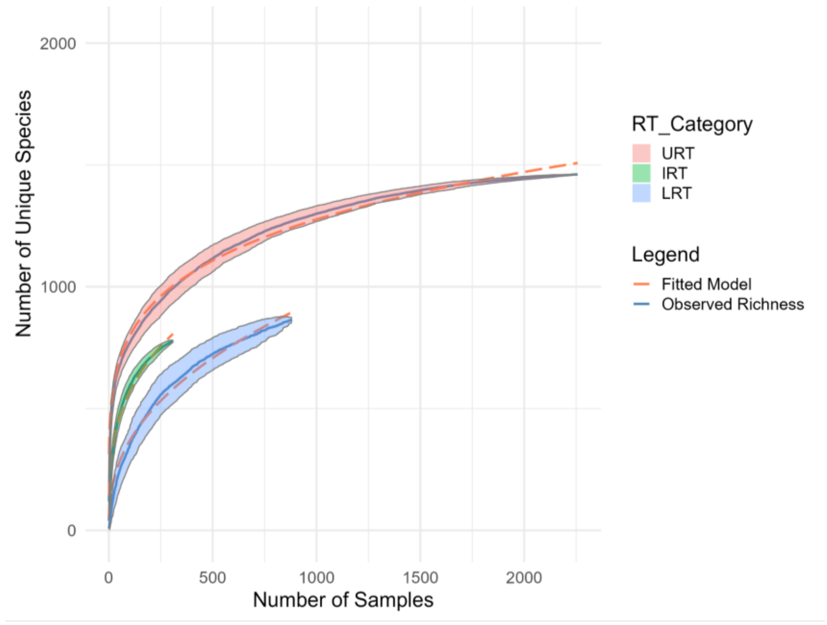


**B.**

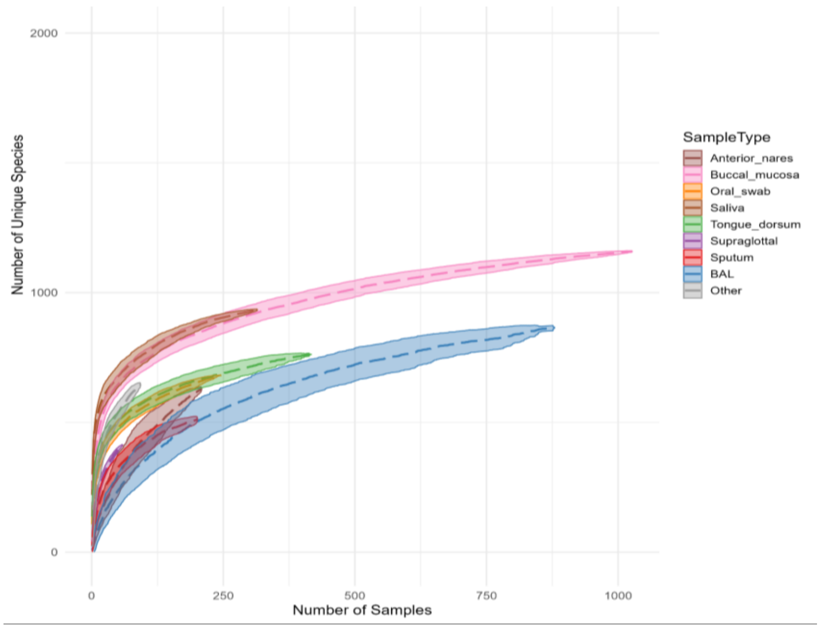


### S-6 Fig.

Species accumulation curves reveal differential saturation across respiratory tract compartments and sample types.

(A) Species accumulation curves (SACs) by respiratory tract category (URT, IRT, LRT) show distinct rates of species discovery with increasing sampling depth. While curves for the intermediate and lower respiratory tract (IRT and LRT) begin to plateau, suggesting partial saturation, the upper respiratory tract (URT) curve continues to rise steadily, indicating incomplete sampling and potential for continued species discovery. (B) SACs by individual sample types show similar trends, with oral sites (e.g., buccal mucosa, saliva, tongue dorsum) yielding more species overall and exhibiting steeper accumulation curves compared to BAL and anterior nares samples. Each curve represents the mean of 1000 random permutations, and shaded regions indicate 95% confidence intervals. Curves were computed on species-level profiles filtered to retain taxa present in ≥10 samples using the specaccum() and fitspecaccum() functions from the vegan R package (v2.6-4), visualized with ggplot2.

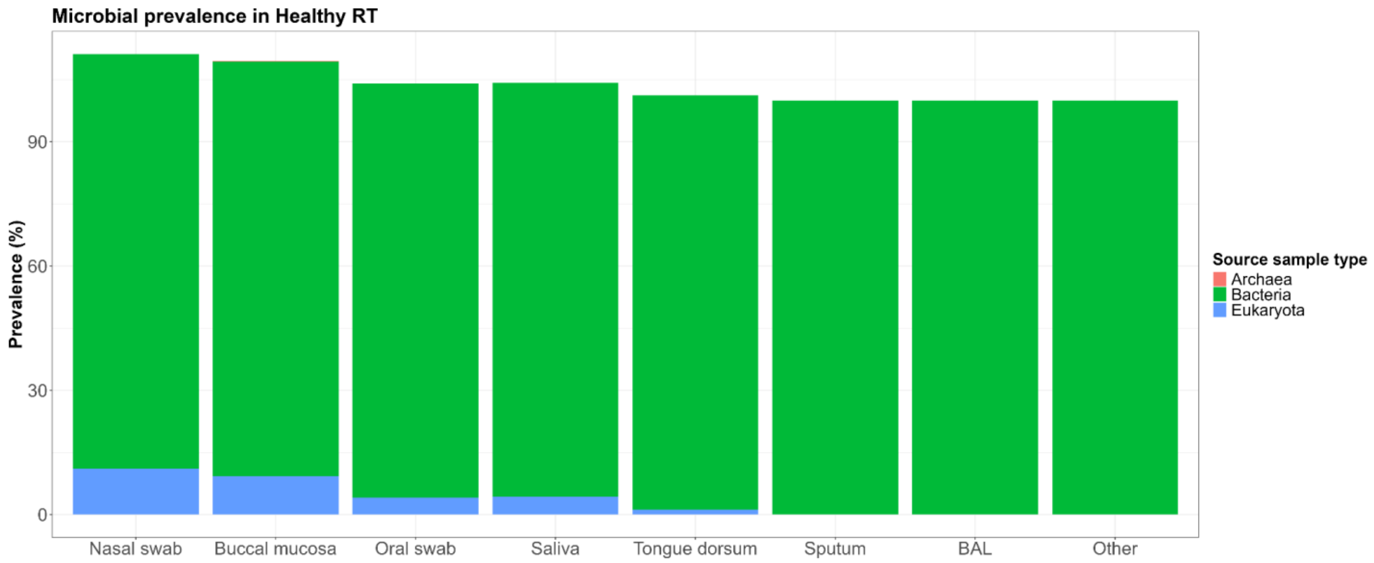


### S-7 Fig.

Microbial domain-level prevalence across respiratory tract sample types in healthy individuals:
Bar plot showing the prevalence of microbial taxa from three major domains of life, viz, Bacteria, Archaea, and Eukaryota across different respiratory tract sample types. Prevalence is defined as the percentage of samples within each sample type in which at least one taxon from the respective domain was detected (abundance > 0). Bacterial taxa were consistently detected in nearly all samples, while Eukaryotic and Archaeal taxa showed limited and sample-type-specific presence.

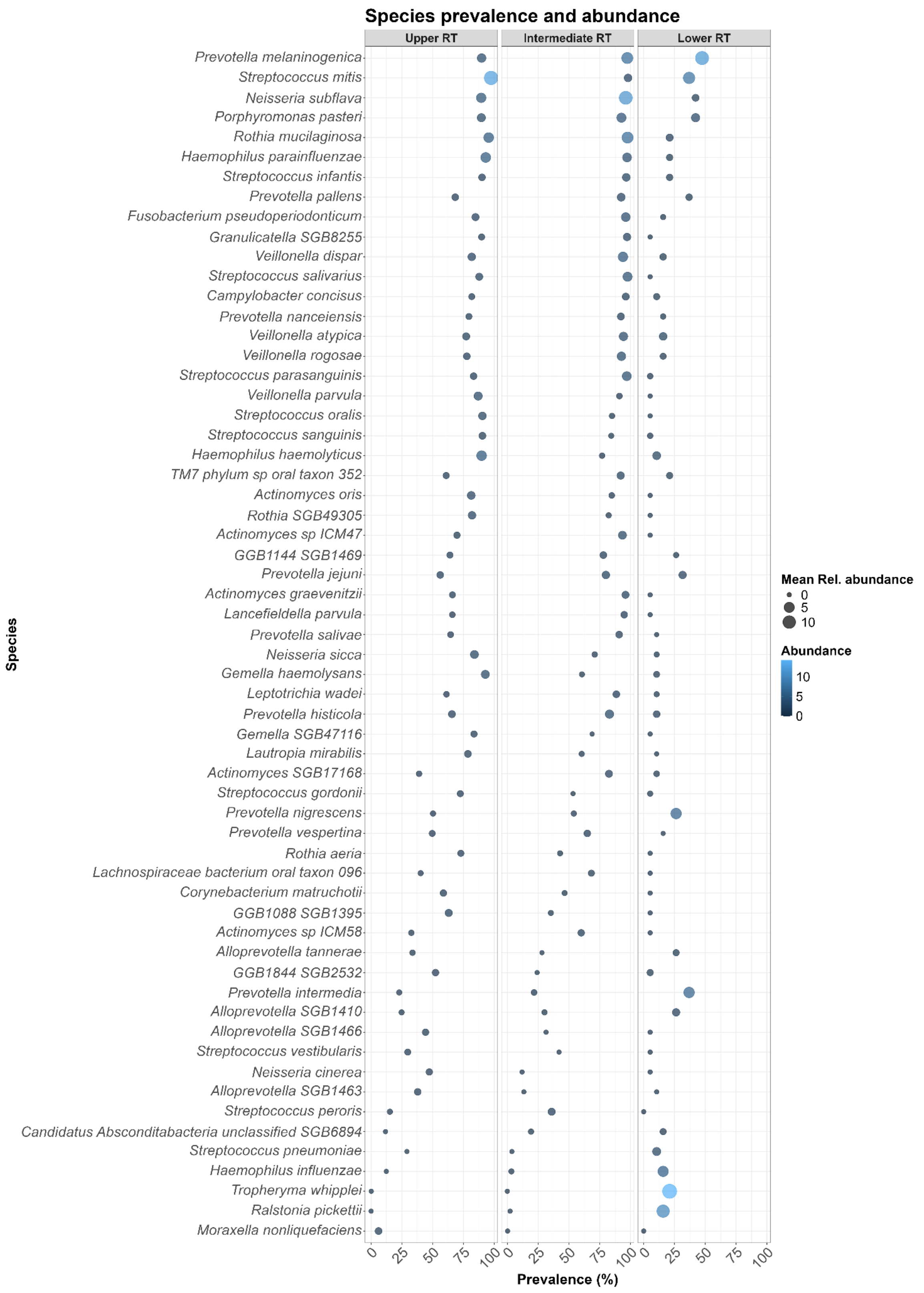


### S-8 Fig.

Taxonomic composition of the healthy human respiratory microbiome across anatomical compartments: Prevalence and mean relative abundance of the top 60 most prevalent species across the upper, intermediate, and lower respiratory tract. While several core commensals (e.g., *Prevotella melaninogenica, Rothia mucilaginosa, Streptococcus mitis*) are consistently present across compartments, others show sharp gradients in prevalence, suggesting spatial filtering or niche specificity. A small number of low-prevalence but occasionally high-abundance species, such as *Moraxella nonliquefaciens* and *Ralstonia pickettii*, were also observed in LRT samples.

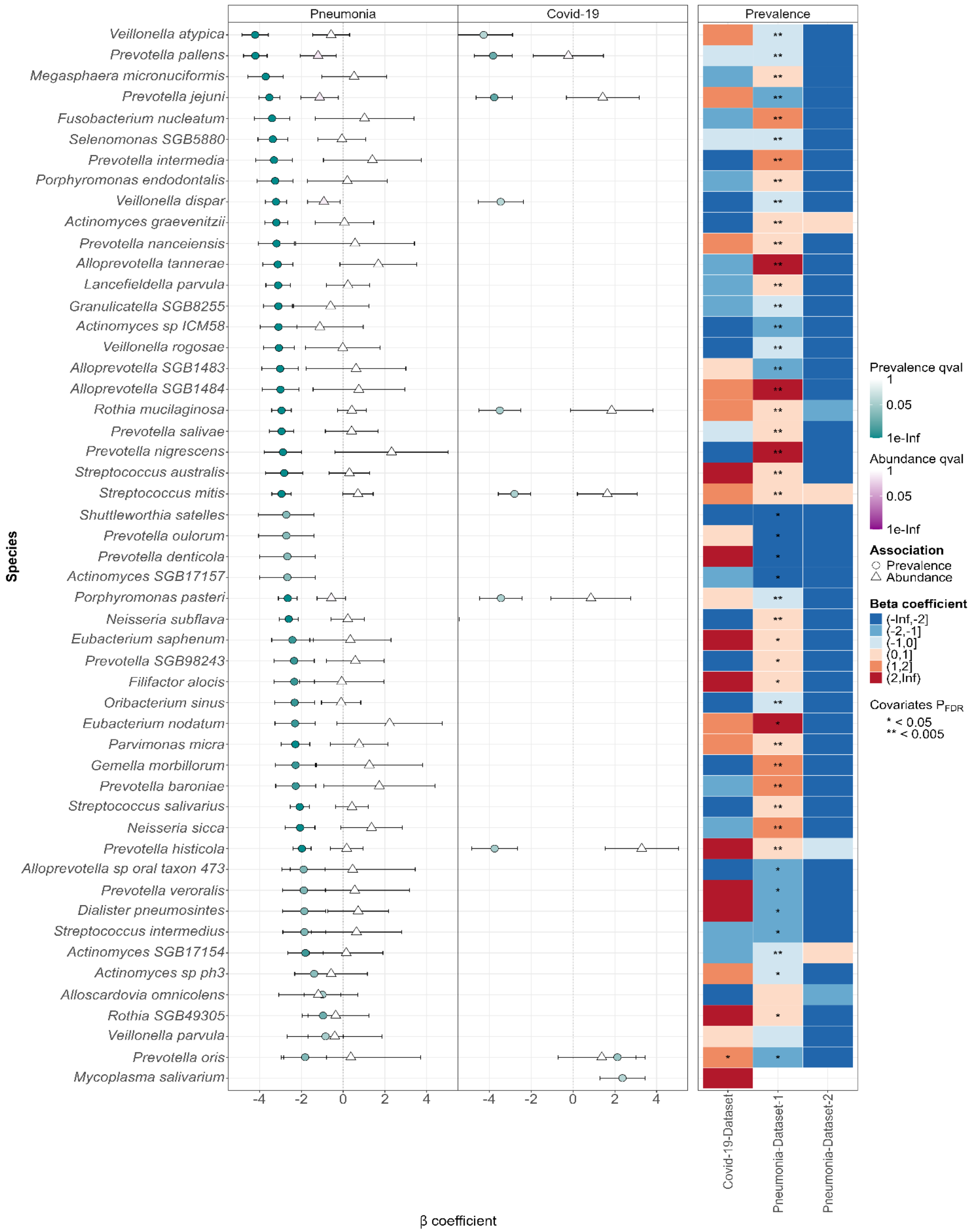


### S-9 Fig.

BAL sample type (LRT): Top 50 disease-associated species from the global model (Model 1), selected based on the q-values (significance, top 50 species with highest significance), excluding uncharacterized GGB/SGB species.

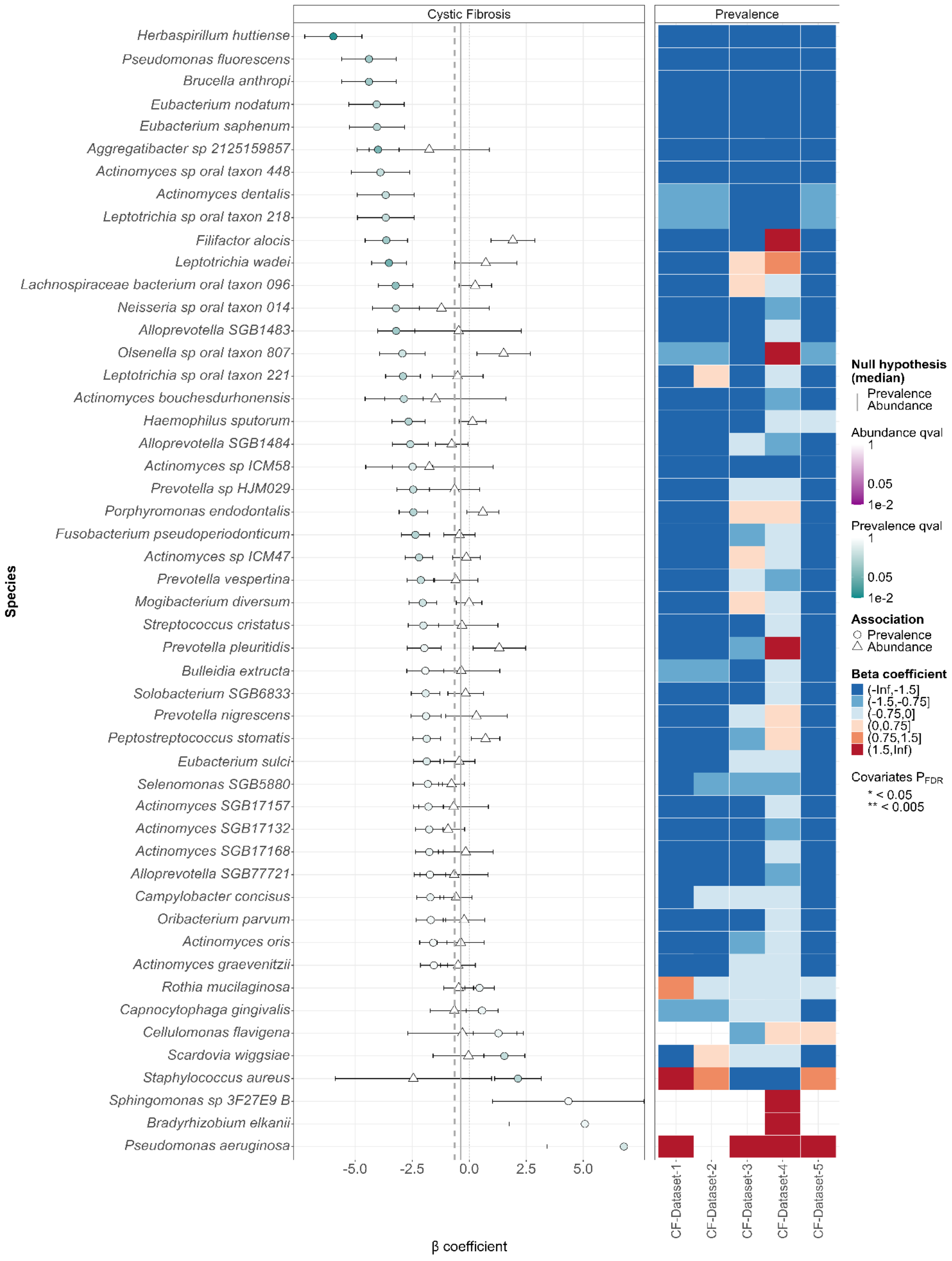


### S-10 Fig.

Sputum sample type (IRT): Top 50 disease-associated species from the global model (Model 1), selected based on the q-values (significance, top 50 species with highest significance), excluding uncharacterized GGB/SGB species.
